## Supplementary Information for "Novel histones and histone variant families in prokaryotes"

#### Supplementary Text

AlphaFold2 and AlphaFold2-multimer are neural networks that predict protein structure given a target sequence [1, 2]. It does so by analyzing coevolutionary information in the form of a multiple sequence alignment (MSA). The MSA is constructed by searching for and aligning sequences that are similar to the target sequence. As a result, the MSA contains evolutionary information on the target sequence. AlphaFold2 participated in the 14th edition of the Critical Assessment of Structure Prediction (CASP) competition. In CASP, research groups test their structure prediction models against a set of proteins. The experimental structures of this set are only known by the organizers of CASP, thus CASP is a good independent way for research groups to test the accuracy of their structure prediction models. AlphaFold2 has demonstrated extremely high accuracy in its ability to correctly predict protein structures in CASP14, largely attributed to its use of full MSAs [3]. Furthermore, in the latest CASP15, all the top contenders use AlphaFold2-based methods [4]. One of the critical features of AlphaFold2 is that

it provides two types of metrics that show how confident AlphaFold2 is in its prediction. These are the predicted local difference distance test (pLDDT) and the predicted aligned error (PAE). These confidence values allow us to interpret the prediction in different ways to assess its quality. A pLDDT value is assigned to every residue and ranges from 0 to 100. It is used as a measure of the local confidence of secondary structures within domains and should not be used to evaluate the relative position of domains. When visualizing AlphaFold2 structures, residues are commonly colored by their pLDDT values, from red (pLDDT = 0) to orange (pLDDT = 50) to yellow (pLDDT = 70) to light blue (pLDDT = 90) to dark blue (pLDDT = 100). Residues with pLDDT $\geqq$ 90 are classified as high confidence, 90 > pLDDT > 70 as confident, 70 > pLDDT > 50 as low confidence, and pLDDT < 50 as very low confidence. The very low confidence regime is generally viewed as a prediction of disorder instead of as regions where AlphaFold2 fails to predict the relevant structures [5, 6]. The other confidence metric, PAE, allows us to identify domains and assess AlphaFold2's confidence in the relative position of domains. It is visualized as a two-dimensional plot with the residue numbers on both the x and y-axis. The error at (x,y) is the expected distance error in Å of residue x relative to residue y when residue y is aligned to the true structure. The distance errors are generally colored from red (30Å) to white (15Å) to blue (0Å), although the AlphaFold2 database colors from white (30Å) to green (0Å). In the case of multimer predictions, the residues of all the chains are appended on the y and x-axis and the different chains are separated by thick black lines. Low distance errors (>10Å) indicate that AlphaFold2 is confident in the relative position of the residues in question. Domains can be identified from the PAE plot by comparing the distance errors between residues from the same chain. Confident interfaces between domains can be identified from the PAE by comparing the distance errors between residues from different chains. For example, the PAE plot in Supplementary Fig. 4c shows distance errors of a dimer prediction. The plot is divided into four parts, with the upper left and lower right parts showing the distance errors between residues from within chain A and chain B respectively. The lower left and upper right parts show the distance errors of residues between chain A and chain B. In both chains A and B we can identify two domains, an N-terminal domain and a C-terminal domain, apparent from the two blue squares in both the upper left and lower right corners of the plot. The C-terminal domains of chains A and B form an interaction interface, apparent from the blue squares in the upper right and lower left corners of the plot.

We have predicted the monomer, dimer, tetramer, and hexamer structures of 5823 prokaryotic histone proteins. To assess the quality of this dataset, we have plotted the distributions of three metrics: MSA depth, pLDDT scores, and interface predicted TM scores (ipTM). MSA depth is an important factor as deep

MSAs contain more evolutionary information and thus give higher-quality predictions. MSAs with at least 100 sequences generally give good predictions [1]. 99% of our predictions have an MSA with 100 or more sequences (Supplementary Fig. 1). Only a small group of histones have shallow MSAs (Supplementary Fig. 1). These are "rare" histones which are part of small categories with few related proteins in the databases that we search to construct the MSAs. For the pLDDT, we show the distribution of the averaged pLDDT scores, meaning that the pLDDT for every residue of a prediction is summed together and divided by the total number of residues (Supplementary Fig. 2). 99%, 95%, 91%, and 86% of monomer, dimer, tetramer, and hexamer predictions respectively have an averaged pLDDT >70, showing that AlphaFold2 is confident in the local structure predictions of the vast majority of histones. The transmembrane histones perform badly on the pLDDT metric due to AlphaFold2 giving low pLDDT values to the transmembrane domains. For every multimer prediction, AlphaFold2 calculates an ipTM score. The ipTM score condenses the information of interactions between chains as visualized in the PAE plots into a single value. The ipTM score ranges from 0 to 1. The ipTM score is high if the majority of residues between chains have low distance errors and is low if the majority have high distance errors. While a high ipTM value (>0.75) is a strong indication of a high-quality multimer prediction, lower ipTM values between 0.3 and 0.75 do not necessarily indicate a low-quality prediction. This is because ipTM is calculated across the whole chain and thus does not reflect if the predicted multimer contains an interface of high quality. For example, bacterial dimer histones and ZZ histones share a highly similar histone fold (Supplementary Fig. 37). ZZ histones, however, contain an additional C-terminal domain which is linked to the histone fold through a disordered linker (Supplementary Fig. 38). The distance errors for the dimer prediction of the bacterial histone Q6MRM1 are all very low and as a result, the ipTM value is very high (0.921) (Supplementary Fig. 39a). For the dimer prediction of the ZZ histone D0LYE7, the predicted dimer interface is identical and the distance errors are also very low, however, the ipTM score is significantly lower (0.466) because the additional N-terminal domain of the ZZ histone does not participate in the multimerization interface (Supplementary Fig. 39b). Thus, while ZZ histones and bacterial histones are predicted to have the same dimerization interface with similarly low distance errors, the ipTMs differ significantly because it is calculated across the whole chain. While some of our multimerization predictions have ipTM scores below 0.75, this does not exclude them from being high-quality multimer predictions. This is the primary reason why we categorized all histones by viewing each prediction manually and assessing the quality of the multimer based on the PAE plot. We provide the PAE plots of every prediction we discuss in Supplementary Fig. 4.

#### Supplementary Tables

Table 1: Data collection and refinement statistics for HTkC. Values for the outer shell are given in parentheses.

| | **HTkC** |
|---|---|
| **Data Collection** | |
| Space group | $P2_12_12_1$ |
| Cell dimensions | |
| $a, b, c$ (Å) | 52.66, 64.29, 79.85 |
| $\alpha, \beta, \gamma$ (°) | 90, 90, 90 |
| Resolution range (Å) | 44.00-1.84 (1.95-1.84) |
| Completeness (%) | 99.6 (99.0) |
| Redundancy | 7.16 (7.15) |
| $< \frac{I}{\sigma(I)} >$ | 12.63 (1.36) |
| $R_{meas}$ | 0.107 (1.48) |
| **Refinement** | |
| No. of reflections, working set | 21711 |
| No. of reflections, test set | 1205 |
| Final $R_{cryst}$ | 0.204 |
| Final $R_{free}$ | 0.237 |
| R.m.s. deviations | |
| Bonds (Å) | 0.005 |
| Angles (°) | 1.263 |

Table 2: A list of the 17 identified prokaryotic histone categories, accompanied by a short description. Categories that have a studied model histone are visualized in bold text.

| Categories | Description |
|---|---|
| **Bacterial dimer** | $\alpha 3$ histones from bacteria that form only dimers and bend DNA. The model bacterial dimer histone is HBb/Bd0055 from *Bdellovibrio bacteriovorus* (PDB: 8FVX,8CMP). |
| Beta | Histones that are predicted to form a unique tetrameric structure. The monomer of these beta histones contains four histone folds on the N-terminus. The C-terminus contains a beta-propeller domain similar to the cartilage acidic protein 1. Likely does not bind DNA. |
| Coiled-coil | Histones that have a coiled-coil domain on their C-terminus. These histones bridge DNA. In the predicted tetramer structure two histone dimers stand opposite each other, interacting through their C-terminal domains, and face away from each other. |
| **DUF1931** | Double histones similar to aq328 from *Aquifex aeolicus* (PDB: 1WWI). |
| Face-to-face | $\alpha 3$ histones that form a characteristic tetramer whereby two dimers stand parallel against each other and directly interact with each other through their histone folds. |
| GTPase | $\alpha 3$ histones that have a eukaryotic-like small Rab GTPase domain on their N-terminus. |
| Halo | Double histones from Halobacteria. These histones form dimer structures which are similar to the nucleosomal tetramer structures. However, based on our AlphaFold predictions, they can not form a superhelix similar to (hyper)nucleosomes. |
| IHF | Histones that have an unknown C-terminal domain, which dimerizes with another tail. The genes of these histones are always found in an operon that also contains an IHF-like protein. |
| **Methanococcales** | Histones that have a C-terminal domain that facilitates DNA bridging. Histone MJ1647 is the model histone of this group (PDB: 8BDK). They are exclusively found in Methanococcales archaea. |
| NCC | Histones that have a coiled-coil domain on their C-terminus. In terms of sequence they are significantly different from coiled-coil histones. They are exclusively found in Nanohaloarchaea. |
| **Nucleosomal** | Histones that form the (hyper)nucleosomal tetramer structure. The model nucleosomal histone is HMfB from *Methanothermus fervidus* (PDB: 1A7W and 5T5K). |
| Phage | $\alpha 3$ histones with a C-terminal domain which facilitates DNA-bridging in the tetramer structure. These histones are found in several Caudovirales metagenomes, and also in some bacterial genomes. |
| Poseidoniia | Double histones found exclusively in Poseidoniia archaea. They contain an unknown C-terminal domain which likely used to be a double histone fold based on structural similarities. |
| RdgC | Histones that have a coiled-coil domain on their C-terminus. The genes of these histones are always found in operons that also contain an RdgC-like protein and an unknown transmembrane protein. |
| Thermoplasmatota | Histones that have a considerably extended $\alpha 2$ helix. They often contain disordered C-terminal tails. The genes of these histones are always found in operons that also contain a Xer-like recombinase. |
| TM | Histones that have a transmembrane domain on their C-terminus. Their $\alpha 1$ helices are often missing. They likely do not bind DNA. |
| ZZ | $\alpha 3$ histones highly similar in their histone fold to the bacterial dimers. However, they contain a ZZ-type zinc finger domain on the N-terminus. |

Table 3: Oligonucleotides used for cloning.

| Name primer | Resulting plasmid | Sequence (5'-3') |
|---|---|---|
| HTkC fragment F | pRD503 | AGGAGATATACATATGGCAGAAATGCTGGTTAAGAGC |
| HTkC fragment R | pRD503 | CTTTGTTAGCAGTTAAACATGACGTGCATACAGGGTTTTACG |
| HTkC vector F | pRD503 | GCATTTCTGCCATATGTATATCTCCTTCTTAAAGTTAAACAAAATTATTTCTAGAGGGG |
| HTkC vector R | pRD503 | ACGTCATGTTTAACTGCTAACAAAGCCCGAAAGG |
| HMfC fragment F | pRD551 | CTTTAAGAAGGAGATATACATATGGAAGAAAAACTGCCGTTTGC |
| HMfC fragment R | pRD551 | CTTTGTTAGCAGTTACAGCTTGCTGGTCAGATCAAATG |
| HMfC vector F | pRD551 | CGGCAGTTTTTCTTCCATATGTATATCTCCTTCTTAAAGTTAAACAAAATTATTTC |
| HMfC vector R | pRD551 | CAGCAAGCTGTAACTGCTAACAAAGCCCG |

Table 4: Plasmids created for this study.

| Name | Backbone | Insert | Resistance | Addgene # |
|---|---|---|---|---|
| pRD503 | pET30b | HTkC (TK1040) | Kanamycin | xxx |
| pRD551 | pET30b | HMfC (Mfer0945) | Kanamycin | xxx |

### Supplementary Figures

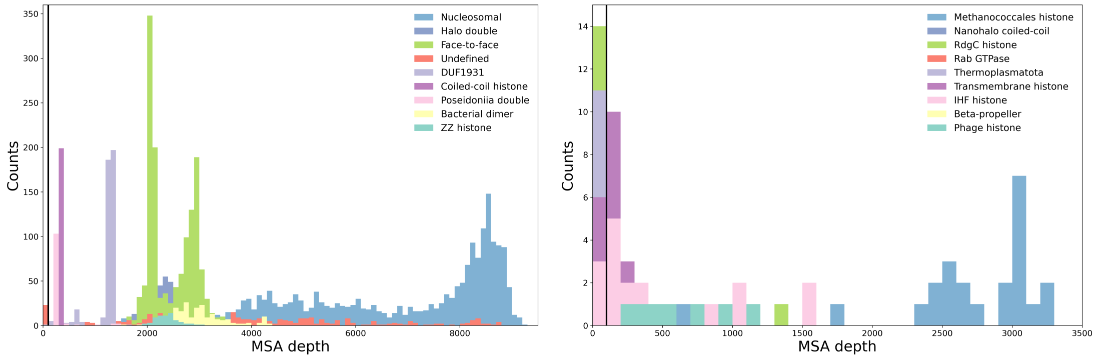

(a) (b)

Figure 1: Histograms showing the distribution of multiple-sequence alignment (MSA) depth of histones from (a) categories with more than 50 members or (b) smaller categories. The vertical black line is placed at an MSA depth value of 100 and depicts the upper boundary for MSAs which we consider to be shallow. Note that the MSA depths of the beta-propeller and Rab GTPase histones are higher than 2500 and thus they fall outside the depicted x-axis range in (b).

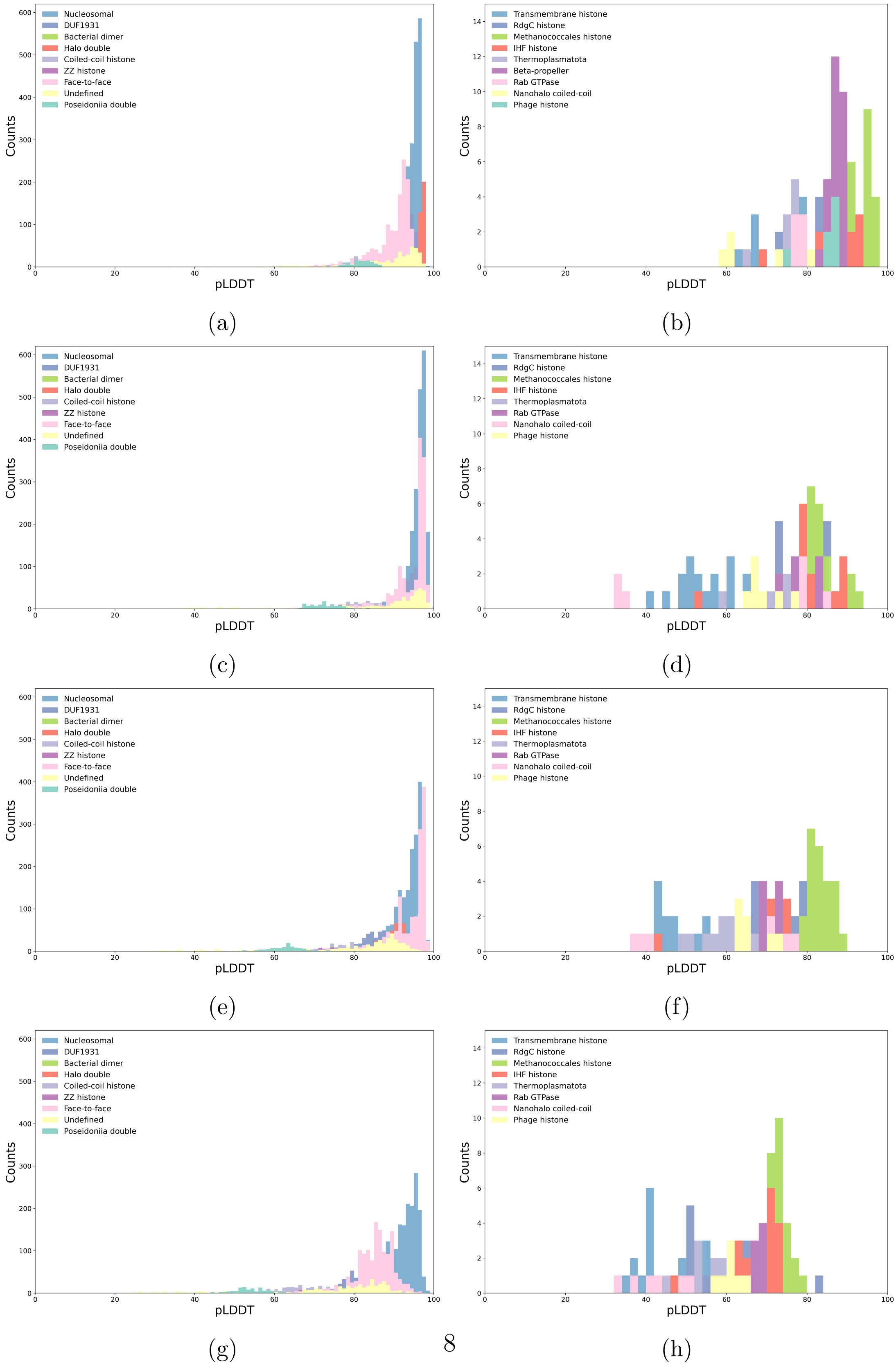

Figure 2: Histograms showing the predicted local distance difference test (pLDDT) values of rank 1 (a,b) monomer, (c,d) dimer, and (e,f) tetramer, (g,h) hexamer predictions from (a,c,e,g) categories with more than 50 members or (b,d,f,h) smaller categories.

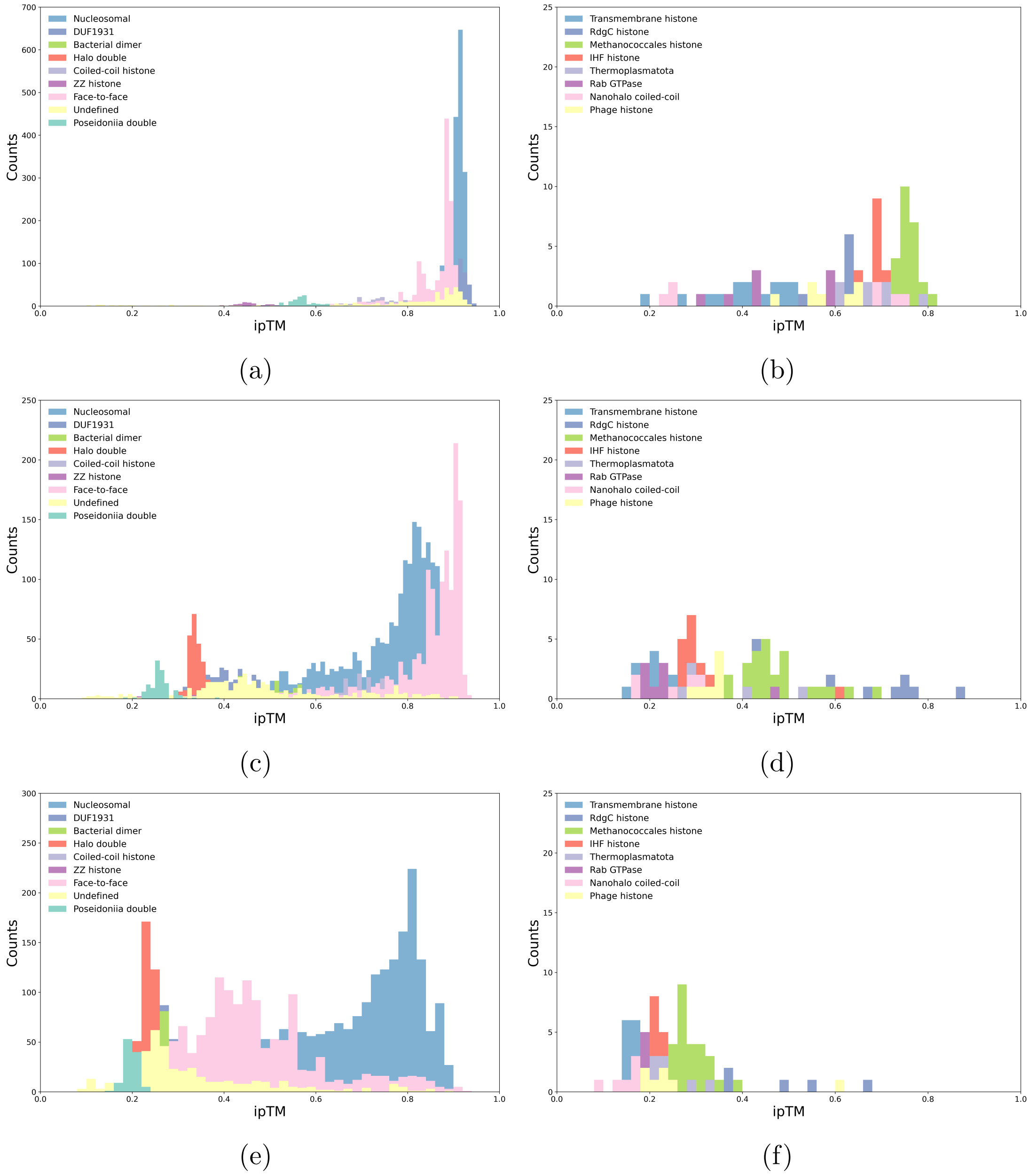

Figure 3: Histograms showing the interface predicted TM score (ipTM) values of rank 1 (a,b) dimer, (c,d) tetramer, and (e,f) hexamer predictions from (a,c,e) categories with more than 50 members or (b,d,f) smaller categories. Note that while high ipTM values (>0.75) likely indicate a high-quality multimer prediction, low ipTM values (<0.75) do not necessarily indicate a low-quality multimer prediction as ipTM is calculated across the whole protein chain.

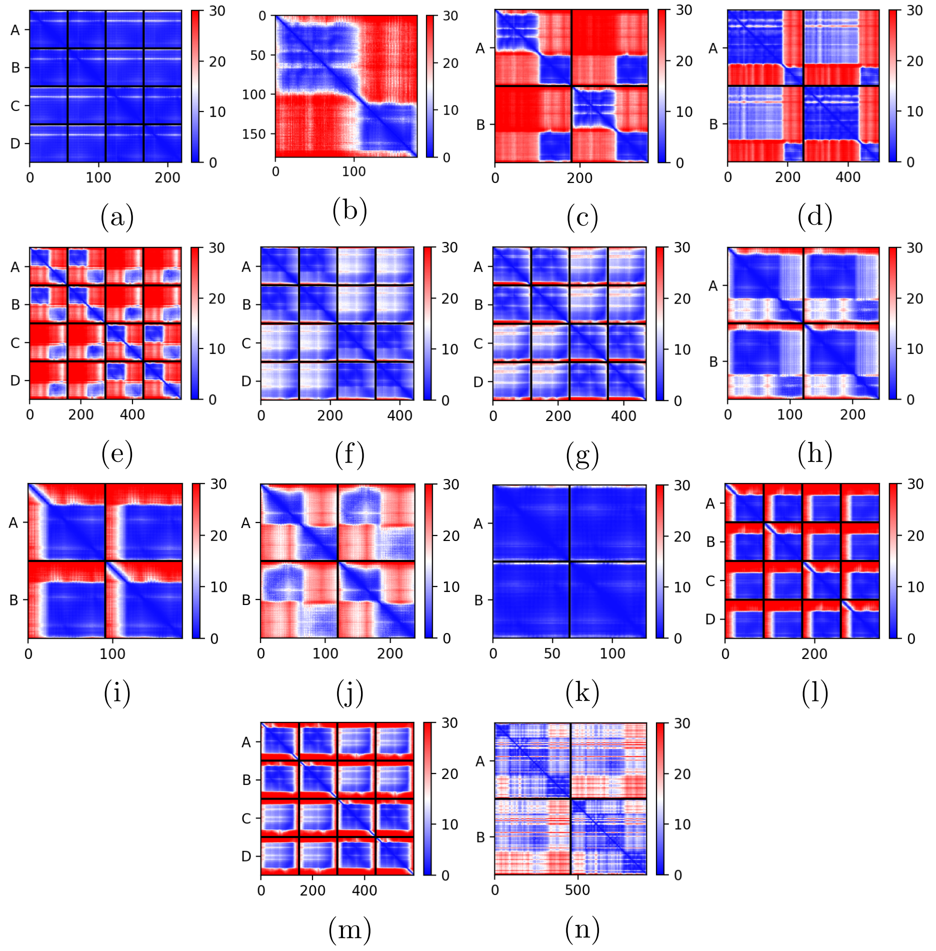

Figure 4: Predicted aligned error plots for (a) D4GZE0 tetramer, (b) D0LYE7 monomer, (c) D0LYE7 dimer, (d) A0A0F8XJF6 dimer, (e) A0A2E7QIQ9 tetramer, (f) E3GZL0 tetramer, (g) D4GVY1 tetramer, (h) A0A358AGI2 dimer, (i) A0A1Q9NJR1 dimer, (j) A0A1F9E2M1 dimer, (k) Q6MRM1 dimer, (l) D0LYZ1 tetramer, (m) A5UK87 tetramer, and (n) Q74P82 dimer. The value at (x,y) is the expected distance error (Å) of residue x relative to residue y when residue y is aligned to the true structure.

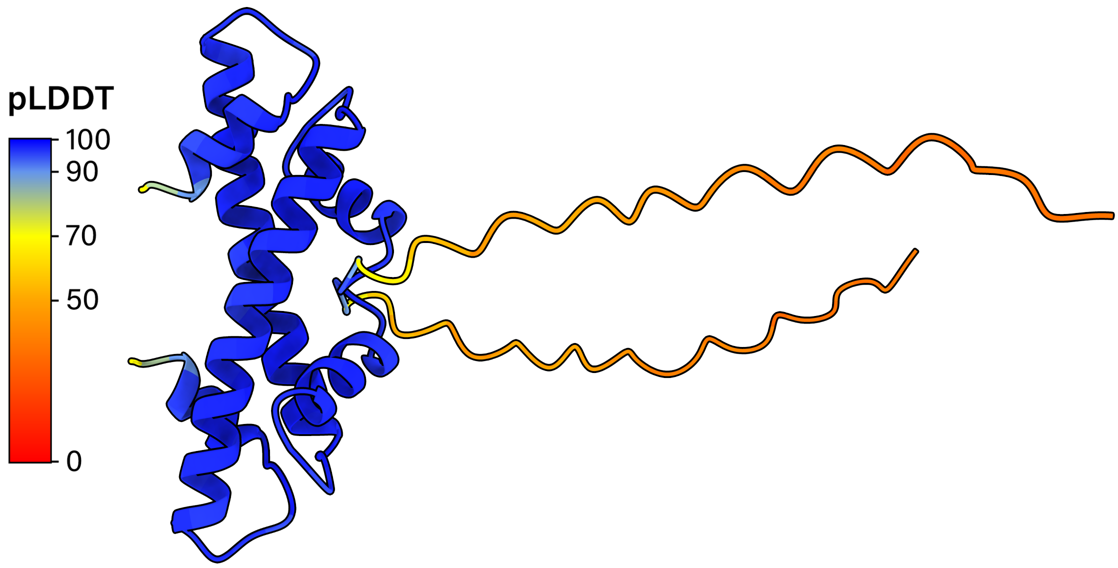

Figure 5: The dimer structure of nucleosomal histone A0A1Q9NJR1 from *Candidatus Heimdallarchaeota archaeon LC3* as predicted by AlphaFold2. Each residue is colored by its pLDDT value. Values below 50 indicate a prediction of disorder.

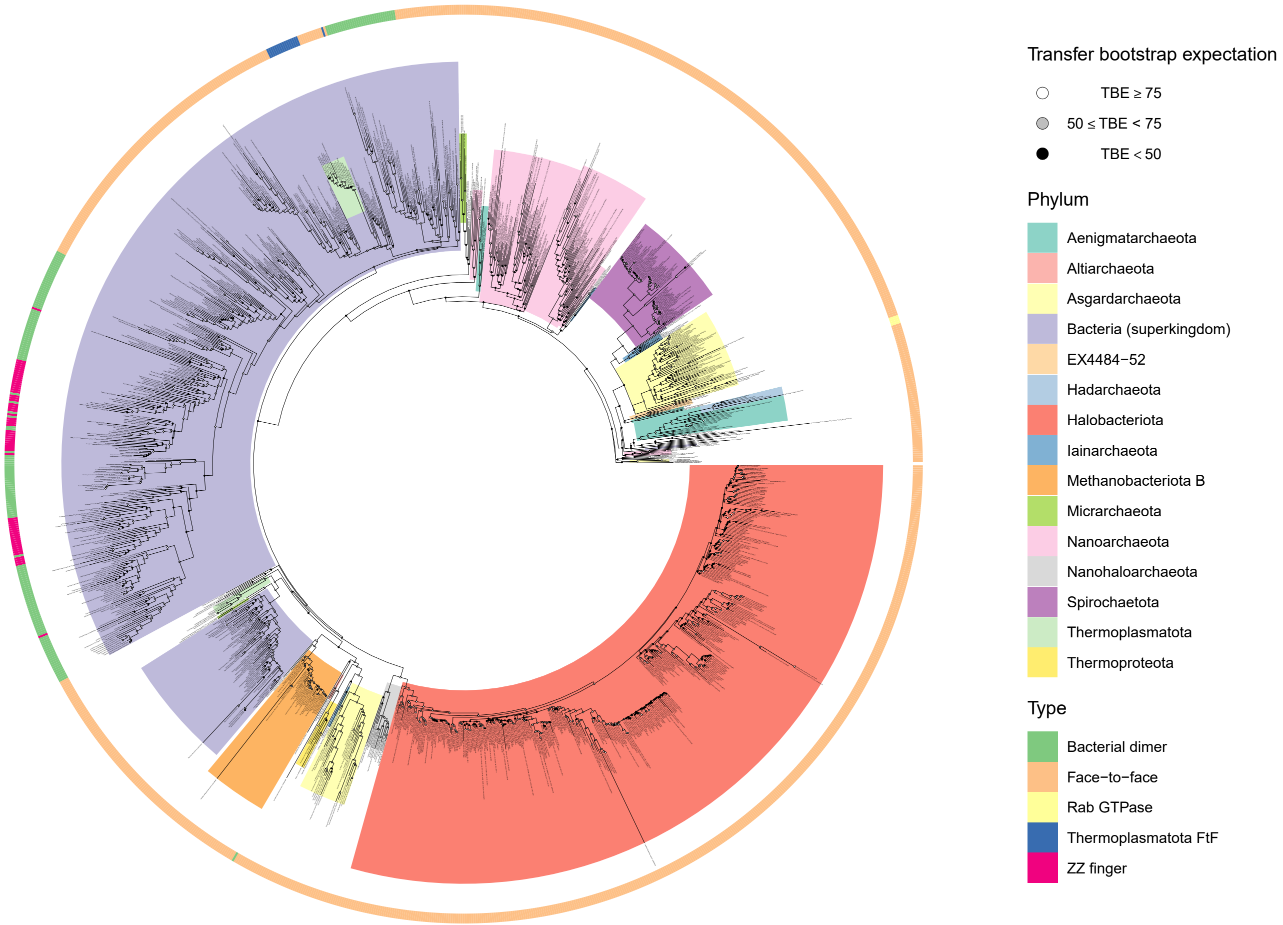
Transfer bootstrap expectation
TBE ≥ 75
50 ≤ TBE < 75
TBE < 50
Phylum
Aenigmatarchaeota
Altiarchaeota
Asgardarchaeota
Bacteria (superkingdom)
EX4484−52
Hadarchaeota
Halobacteriota
Iainarchaeota
Methanobacteriota B
Micrarchaeota
Nanoarchaeota
Nanohaloarchaeota
Spirochaetota
Thermoplasmatota
Thermoproteota
Type
Bacterial dimer
Face−to−face
Rab GTPase
Thermoplasmatota FtF
ZZ finger

(a)

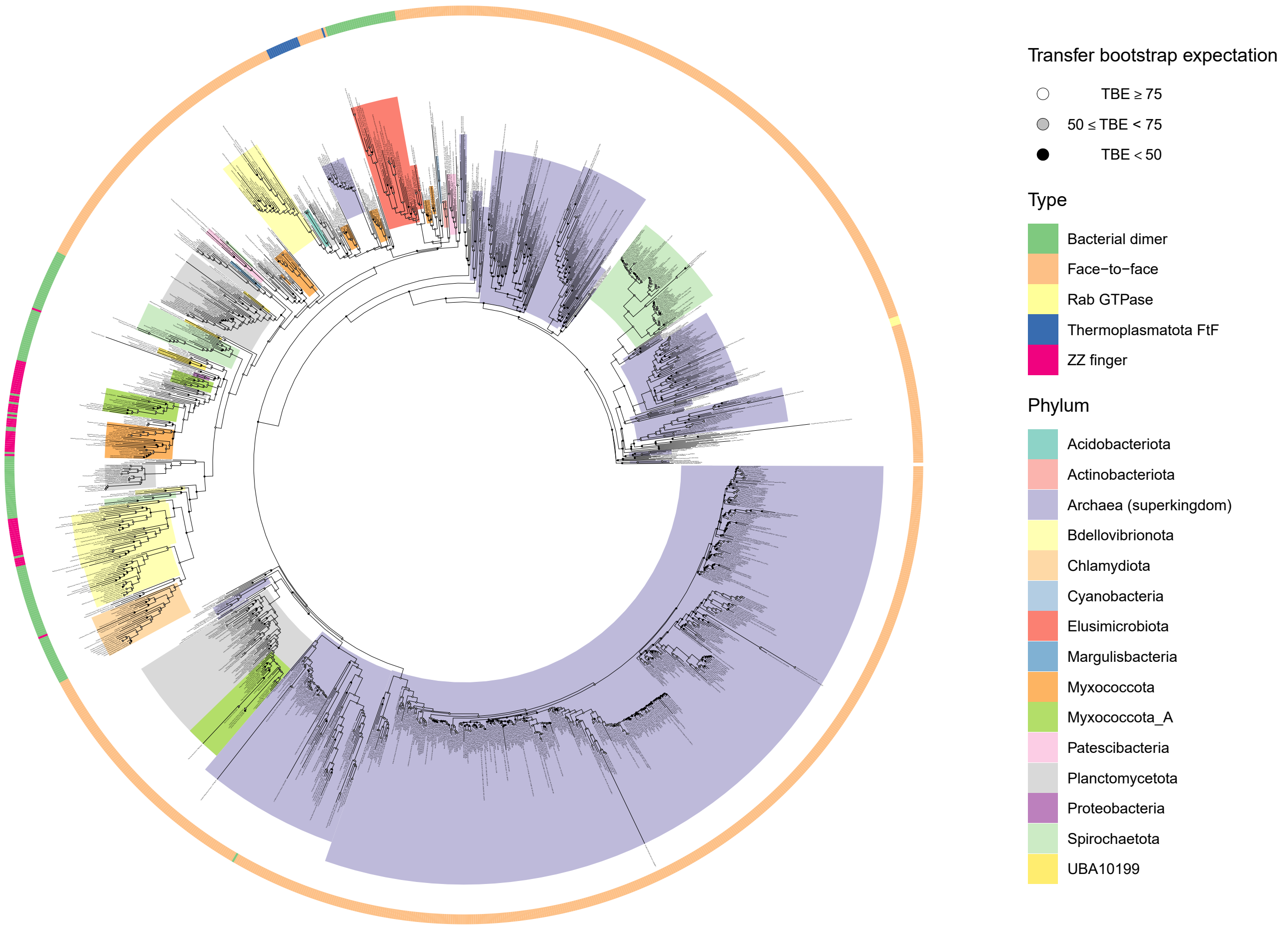

(b)

Figure 6: Phylogenetic tree of face-to-face, bacterial dimer, ZZ, and Rab GTPase histones. The (a) archaeal and (b) bacterial clades are colored by phylum as they are assigned in the GTDB database (v207). The outer ring indicates to which histone category the histone in question belongs. The tree was generated with RAxML-NG. 600 bootstraps were performed and used to calculate the transfer bootstrap expectation values (TBE).

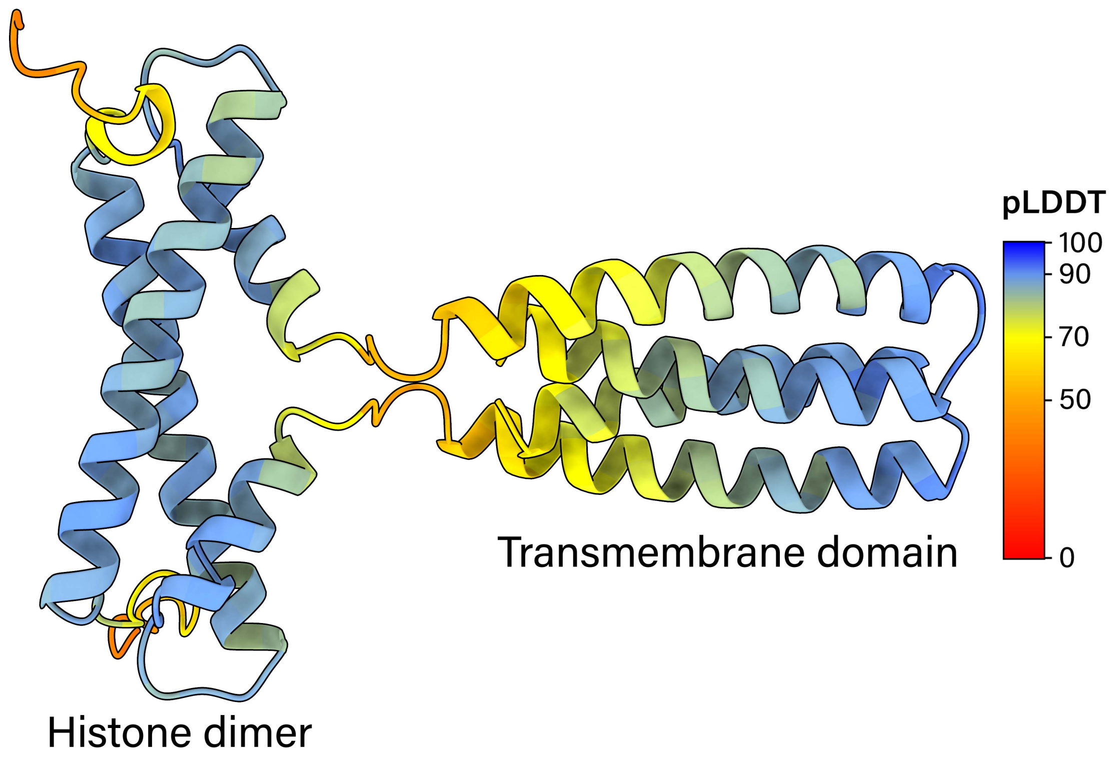

(a)

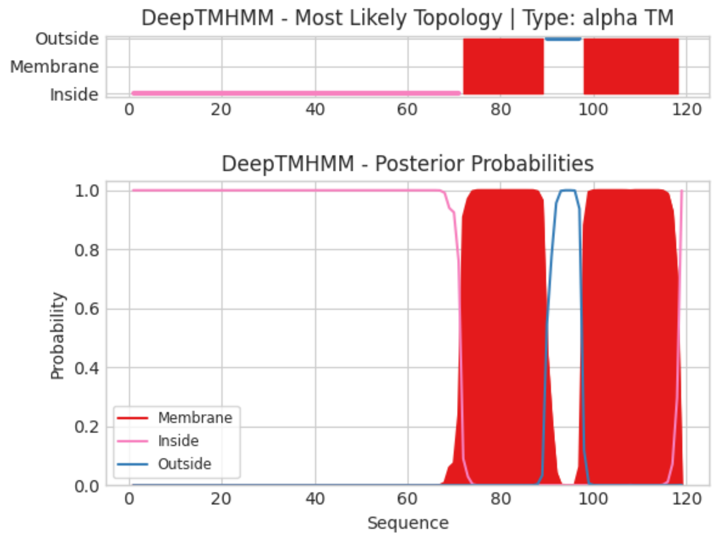

(b)

Figure 7: (a) The dimer structure of transmembrane histone A0A1F9E2M1 as predicted by AlphaFold2. Each residue is colored by its pLDDT value. (b) Prediction of A0A1F9E2M1's transmembrane probability and its topology by DeepTMHMM.

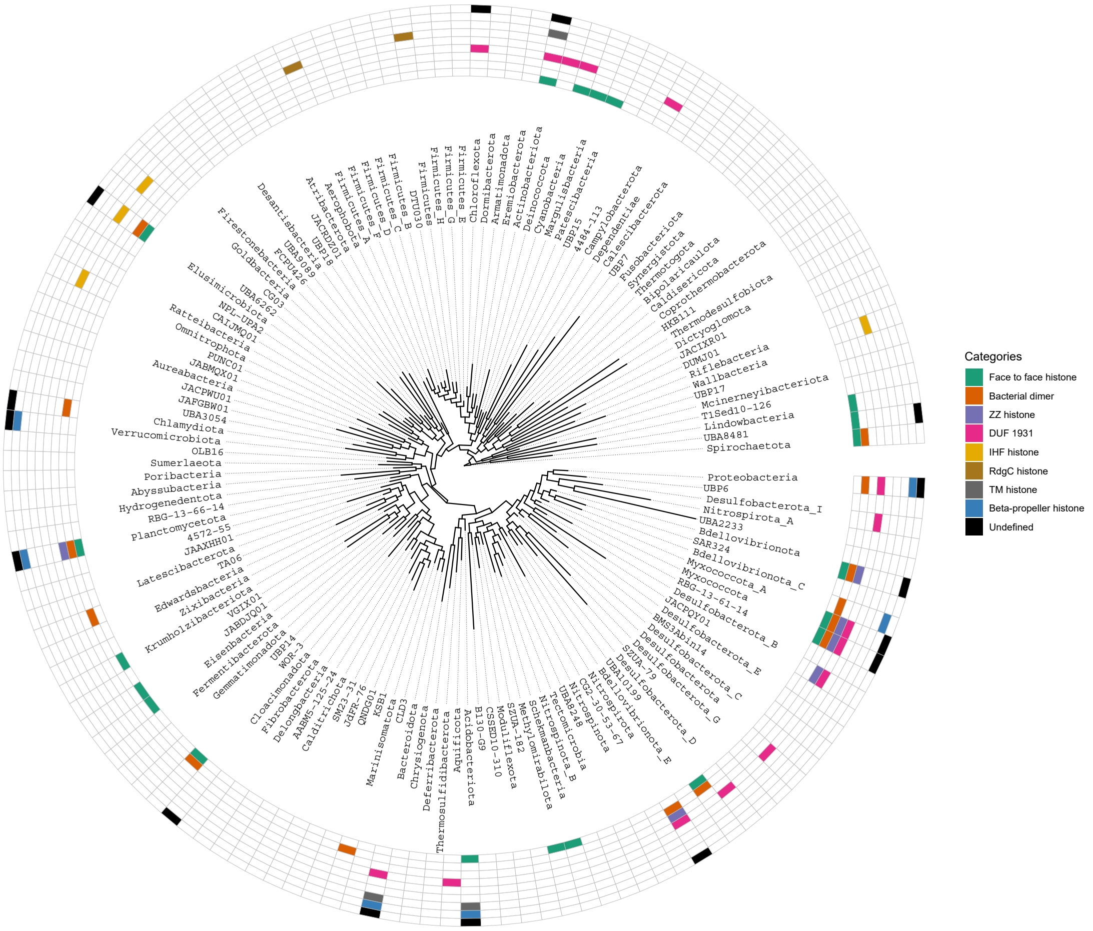

Figure 8: Cladogram of the bacteria superkingdom showing the distribution of face-to-face, bacterial dimer, ZZ, DUF1931, IHF, RdgC, TM, beta-propeller, and undefined histones. The cladogram is based on GTDB version 207.

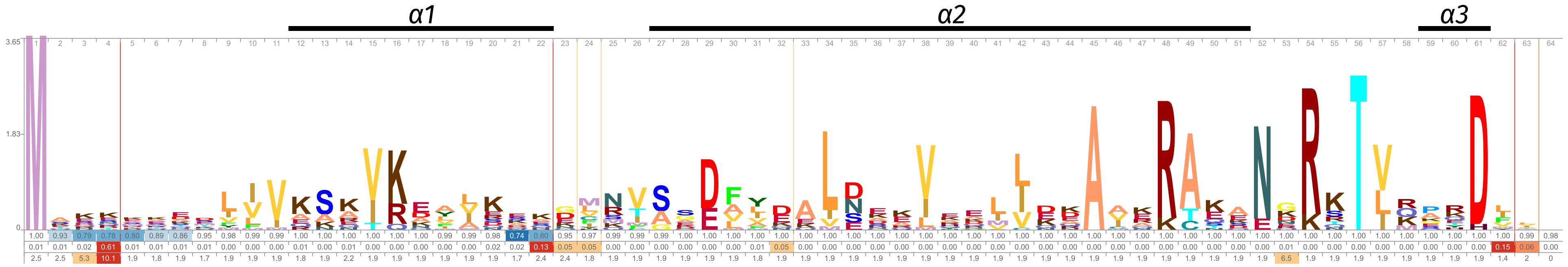

Figure 9: Logo representation of the face-to-face histone HMM profile. Only residues with scores above background frequency are shown. The occupancy probability, insert probability, and insert length values are below the residues. The secondary structure is visualized above the residues.

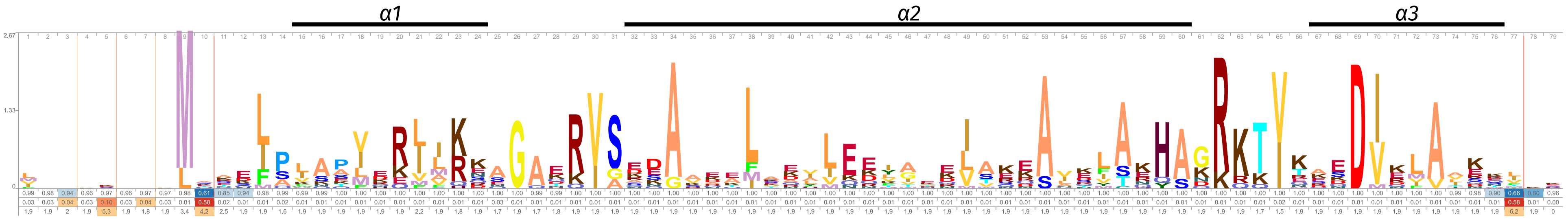

Figure 10: Logo representation of the nucleosomal histone HMM profile. Only residues with scores above background frequency are shown. The occupancy probability, insert probability, and insert length values are below the residues. The secondary structure is visualized above the residues.

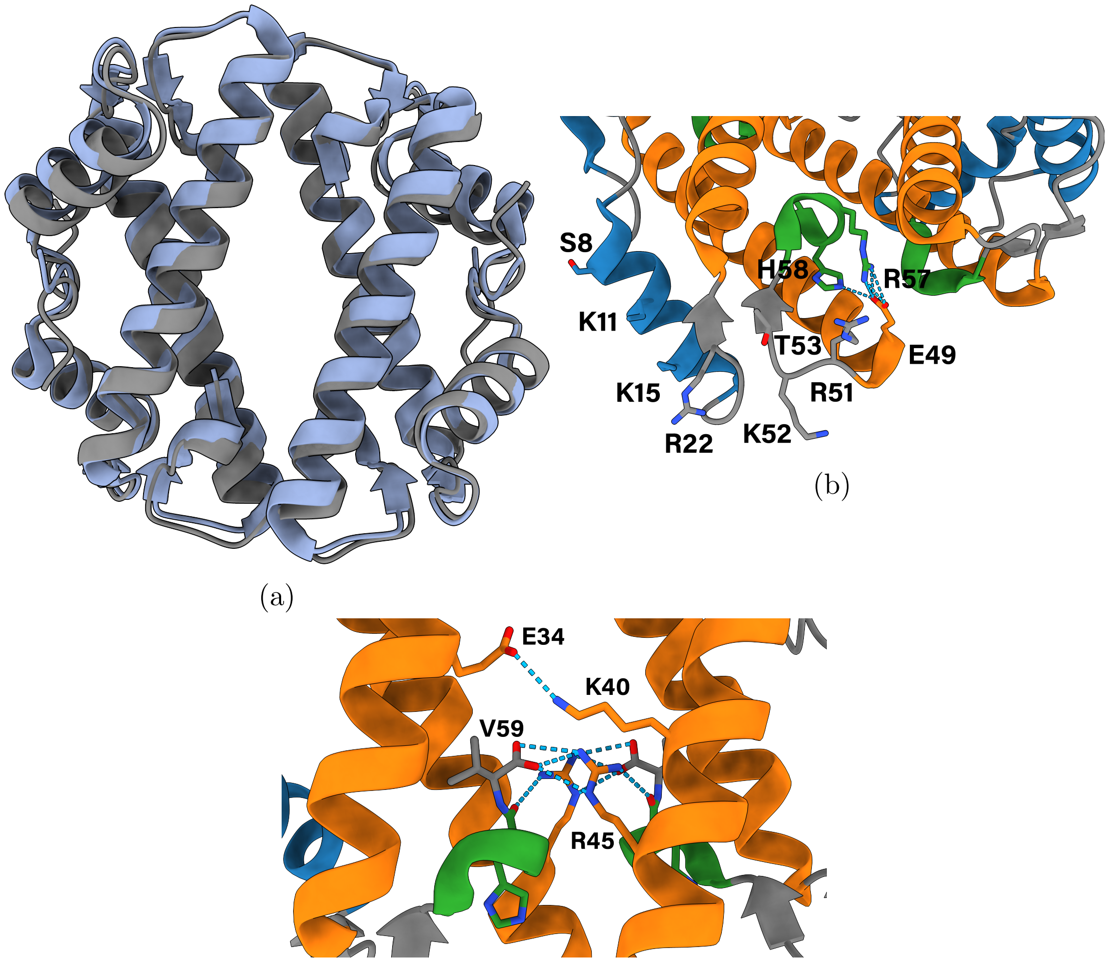

(a)

(b)

(c)

Figure 11: (a) The crystal structure of HTkC (blue) with its AlphaFold prediction (gray). (b) Magnified view of the dyad area of the HTkC crystal, highlighting possible DNA binding residues. (c) Magnified view of the dimer-dimer interface in the HTkC crystal, highlighting R45 which forms salt bridges with the carboxyl terminus of the opposing dimer. An additional salt bridge between the two dimers is made by K40 and E34, which are not conserved in FtF histones outside of the *Thermococci* class.

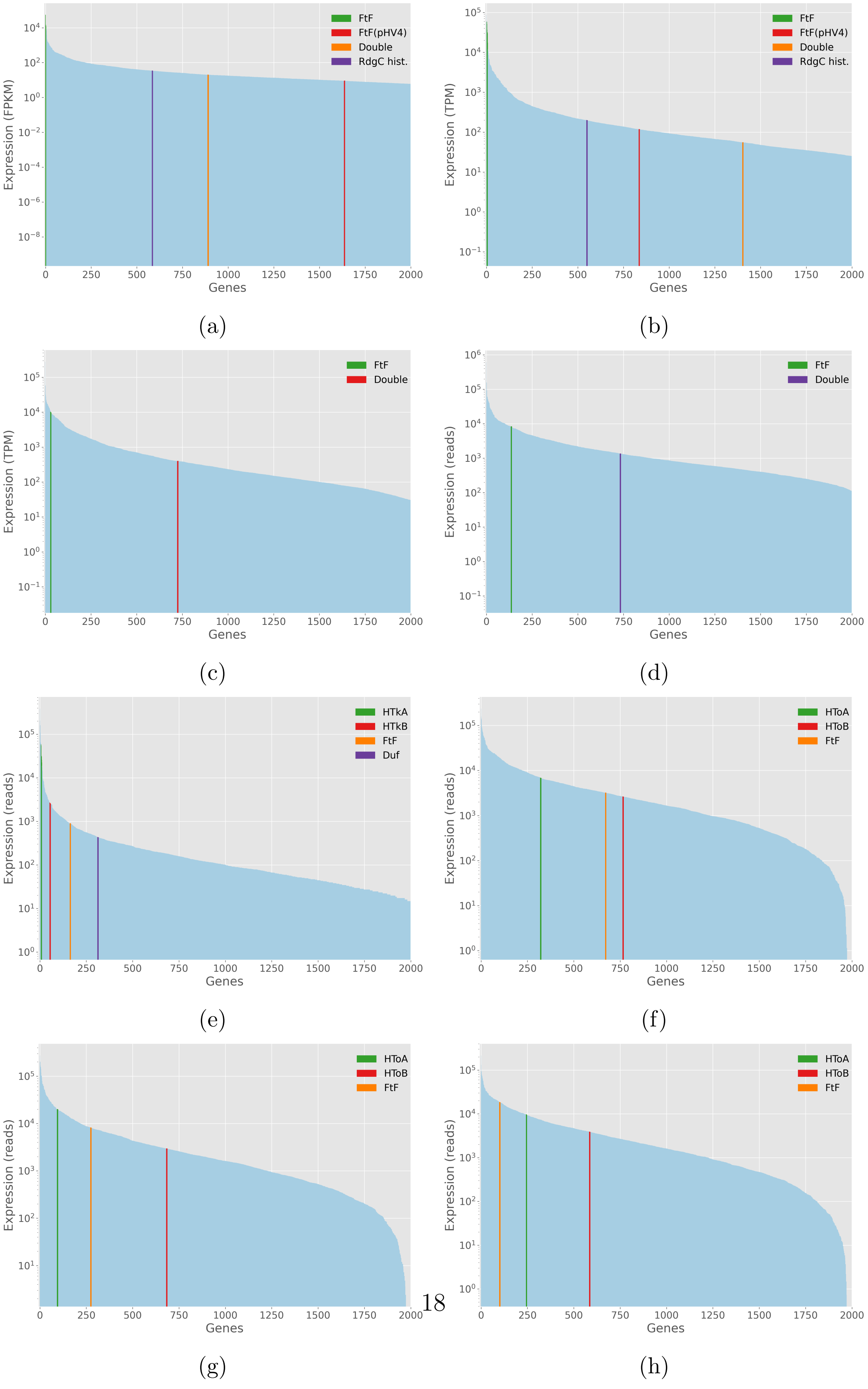
FtF
FtF(pHV4)
Double
RdgC hist.
Expression (FPKM)
Genes
(a)
FtF
FtF(pHV4)
Double
RdgC hist.
Expression (TPM)
Genes
(b)
FtF
Double
Expression (TPM)
Genes
(c)
FtF
Double
Expression (reads)
Genes
(d)
HTkA
HTkB
FtF
Duf
Expression (reads)
Genes
(e)
HToA
HToB
FtF
Expression (reads)
Genes
(f)
HToA
HToB
FtF
Expression (reads)
Genes
(g)
HToA
HToB
FtF
Expression (reads)
Genes
(h)

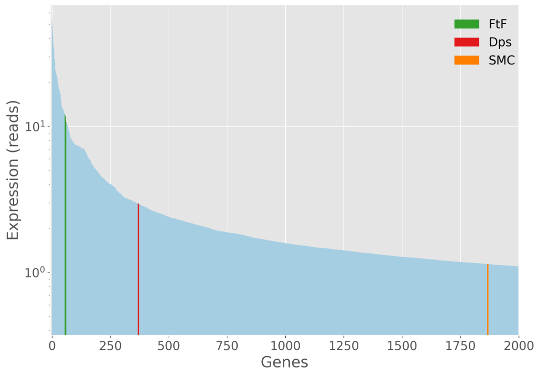

(i)

Figure 12: Transcriptome plots of various archaea and bacteria. Genes are ranked from highest expression (at x=0) to lowest expression. Only the top 2000 expressed genes are visualized. The y-axis is on a base 10 logarithmic scale. Legend abbreviations are FtF:face-to-face histone, FtF(x): face-to-face histone from plasmid x, Double:Halobacterium double histone, Duf:DUF1931, HTkA and HTkB:nucleosomal histones, HToA and HToB:nucleosomal histones, RdgC hist.:RdgC histone. (a) mRNA transcriptome data for *Haloferax volcanii* from Ammar et al.. (b) mRNA transcriptome data for *Haloferax volcanii* from Blombach et al.. (c) mRNA transcriptome data for *Halobacterium salinarum* NRC-1 from Lopez et al.. (d) mRNA transcriptome data for *Halobacterium salinarum* NRC-1 from Sakrikar et al.. (e) mRNA transcriptome data for *Thermococcus kodakarensis* from Jager et al.. (f-h) mRNA transcriptome data for *Thermococcus onnurineus* NA1 in yeast extract-peptone-sulfur (f), modified minimal-CO (g), and modified minimal-formate (h) media from Cho et al.. (i) mRNA transcriptome data for *Leptospira interrogans* serogroup Icterohaemorrhagiae serovar Lai (strain 56601) from Xue et al..

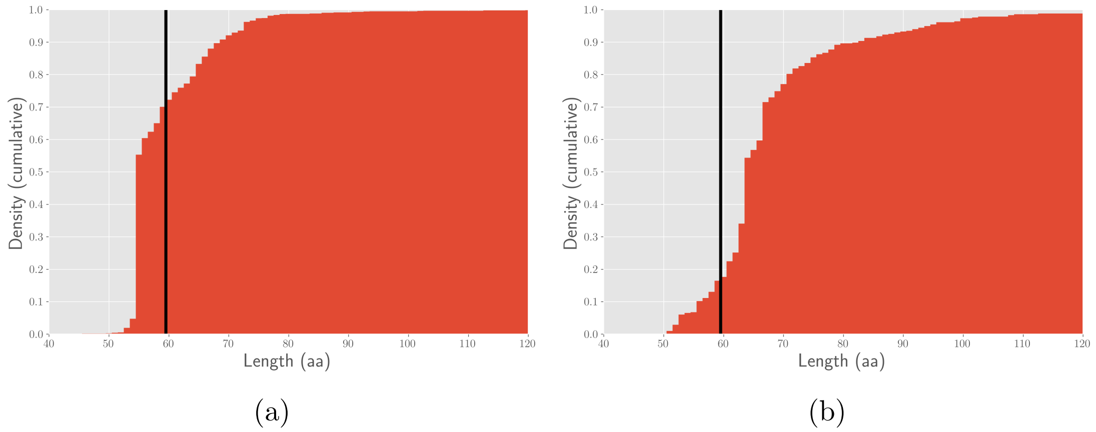

Figure 13: Cumulative density histograms of the sequence length of archaeal (a) and bacterial (b) face-to-face histones. The black line indicates the length of a face-to-face histone without tails.

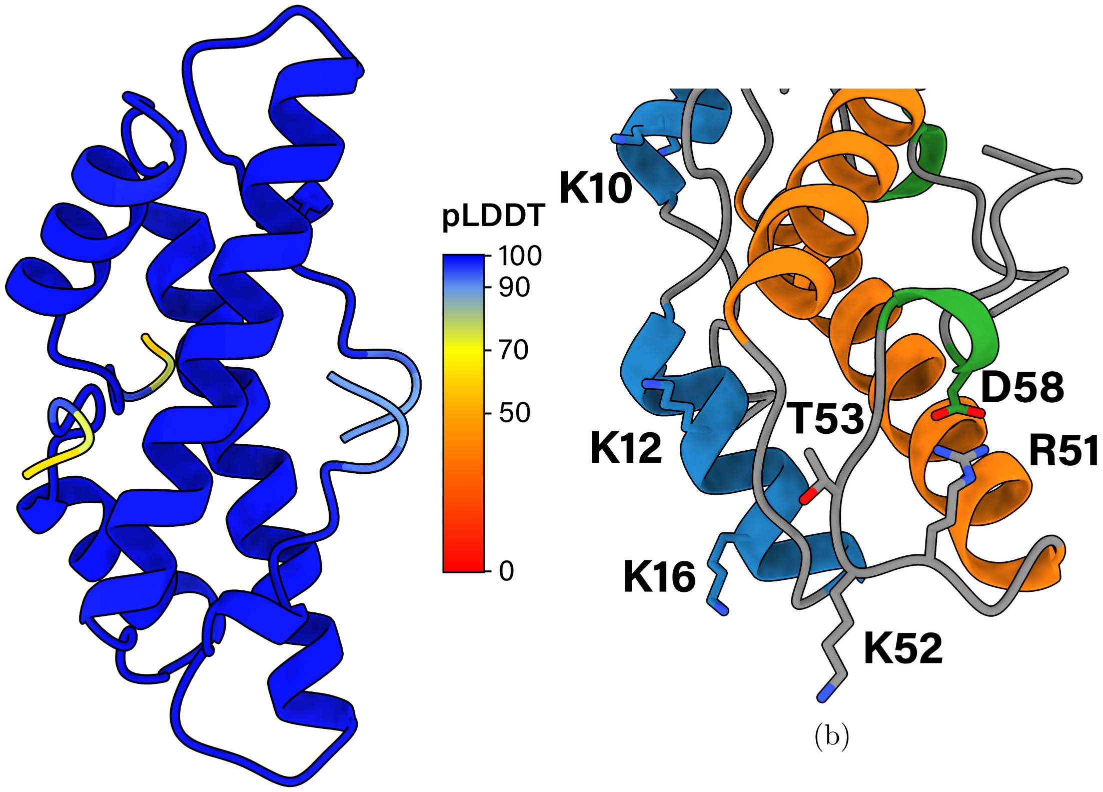

(a) (b)

Figure 14: (a) The homodimer of bacterial dimer histone Q6MRM1 from *Bdellovibrio bacteriovorus* HD100 as predicted by AlphaFold2. Each residue is colored by its pLDDT value. (b) The DNA binding residues K10, K12, K16 and the RKTxxxxD motif of bacterial dimer histone Q6MRM1. Residues K10, K12, K16, R51, K52, T53, and D58 correspond to K11, K13, K17, R52, K53, T54, and D59 in the HMM profile (Supplementary Fig. 15).

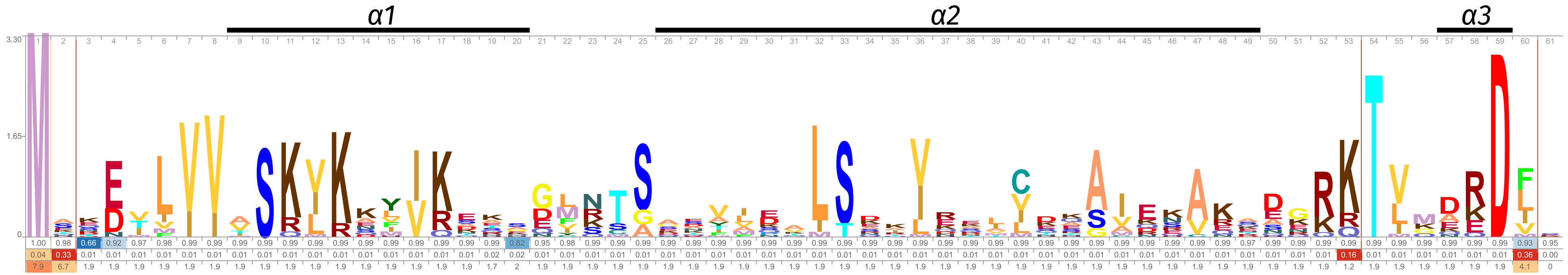

Figure 15: Logo representation of the bacterial dimer histone HMM profile. Only residues with scores above background frequency are shown. The occupancy probability, insert probability, and insert length values are below the residues. The secondary structure is visualized above the residues.

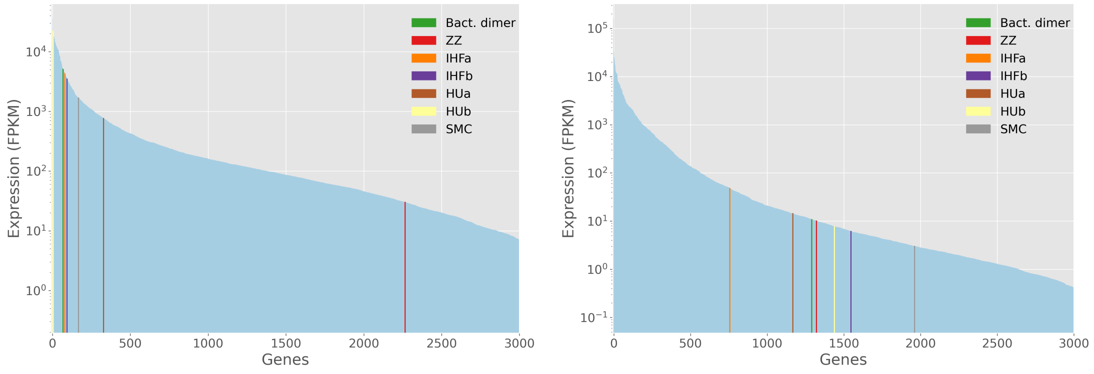

(a) (b)

Figure 16: Transcriptome plots of *Bdellovibrio bacteriovorus* HD100. Genes are ranked from highest expression (at x=0) to lowest expression. Only the top 3000 expressed genes are visualized. The y-axis is on a base 10 logarithmic scale. (a) mRNA transcriptome data for *Bdellovibrio bacteriovorus* HD100 during its growth phase from Karunker et al.. (b) mRNA transcriptome data for *Bdellovibrio bacteriovorus* HD100 during its attack phase phase from Karunker et al.. Cells were grown in the presence of *Escherichia coli* prey MG1655.

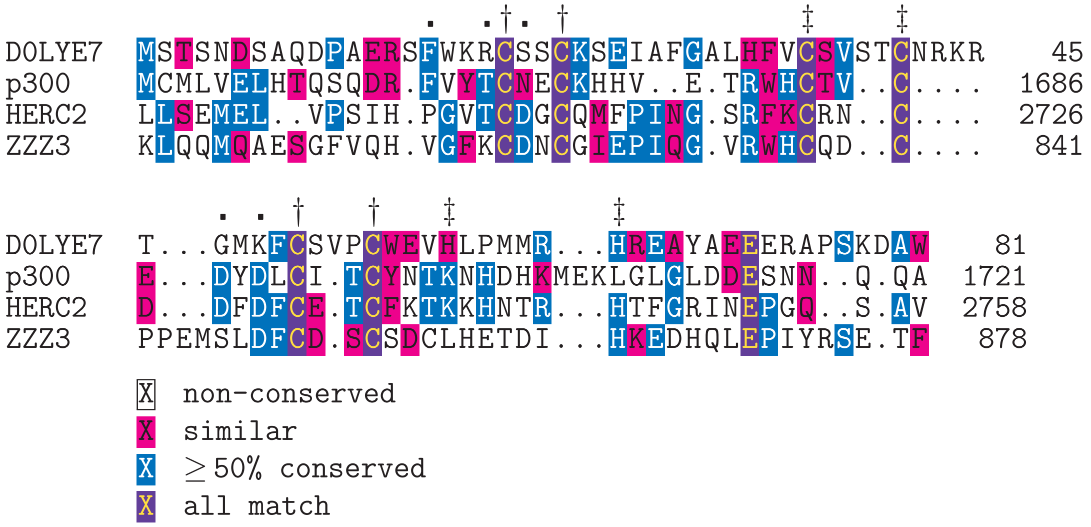

Figure 17: Multiple sequence alignment of the ZZ domains of D0LYE7, p300, HERC2, and ZZZ3. Single daggers (†) mark the C4 zinc motif residues in D0LYE7. Double daggers (‡) mark the C2H2 zinc motif residues in D0LYE7. Dots (.) mark residues of HERC2 that bind the tail of H3.

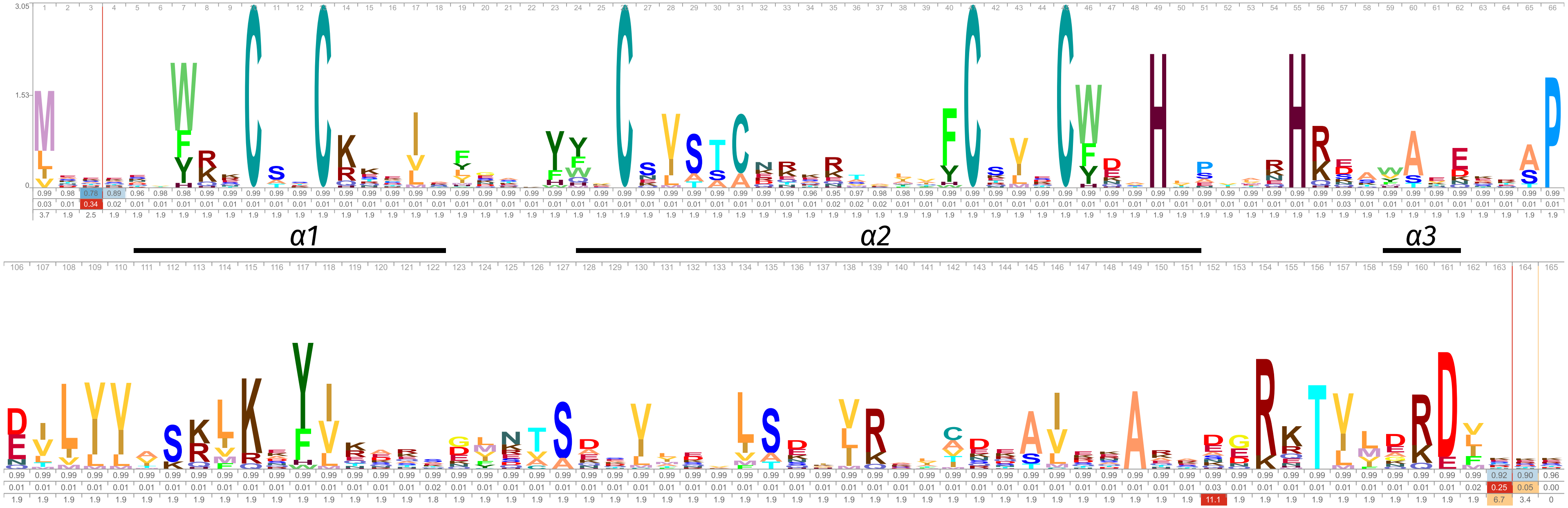

Figure 18: Logo representation of the ZZ histone HMM profile. Only residues with scores above background frequency are shown. The occupancy probability, insert probability, and insert length values are below the residues. The secondary structure is visualized above the residues.

```
H3        MARTK.QTARKSTGGKAPRKQLATKA    25
D0LYZ1    MATKKKKTTKKKAASKTSKKPAAKKE    26
E1WYC3    MAKKK.ATKKAATKKKATKTTKASKK    25
```

```
X  non-conserved
X  similar
X  ≥ 50% conserved
X  all match
```

Figure 19: Multiple sequence alignment of the N-terminal tails of histones H3, D0LYZ1 from *Haliangium ochraceum SMP-2*, and E1WYC3 from *Halobacteriovorax marinus SJ*. Both organisms also contain a ZZ-histone (D0LYE7 and E1WXM4 respectively). The alignment was made with TCoffee using default parameters.

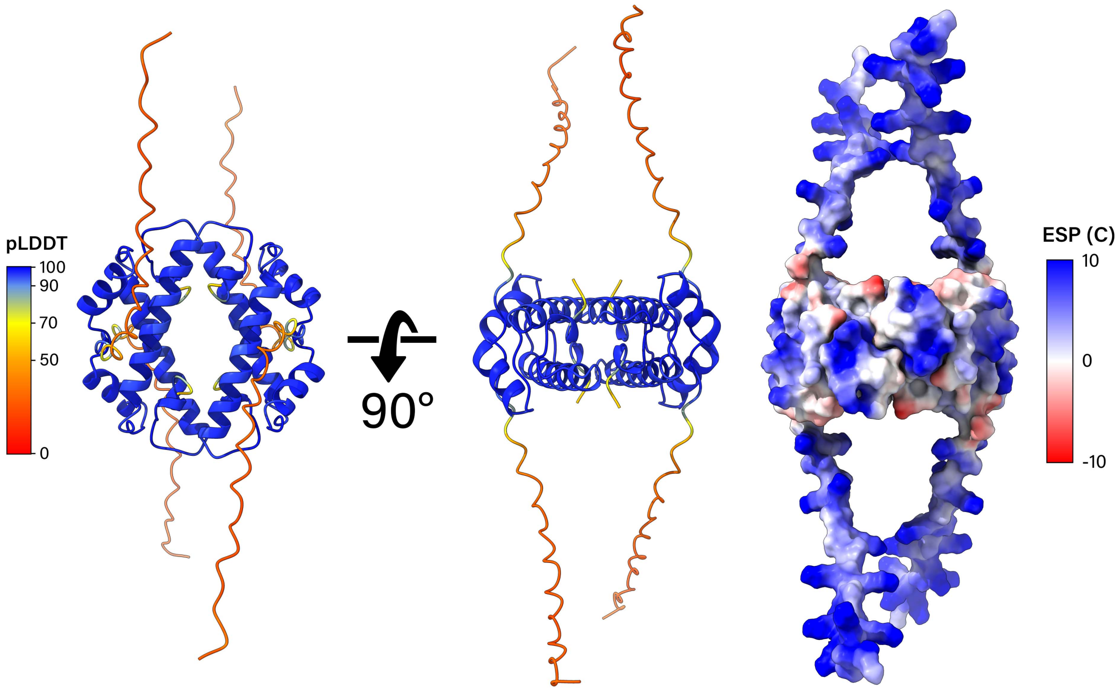

Figure 20: (a) The homotetramer of D0LYZ1 as predicted by AlphaFold2. D0LYZ1 is a face-to-face histone from *Haliangium ochraceum SMP-2*. The first two structures from the left are colored by their pLDDT value for each residue. Values above 70 indicate confidence in the local structure. Values below 50 indicate a prediction of disorder. The rightmost structure visualizes the molecular surface of D0LYZ1 and is colored by Coulombic electrostatic potential (ESP).

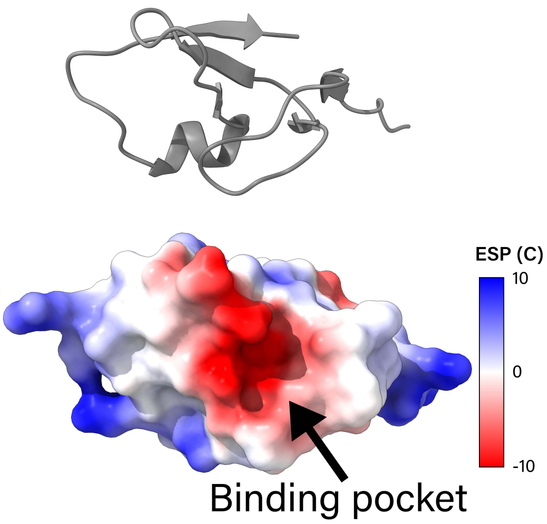

(a)

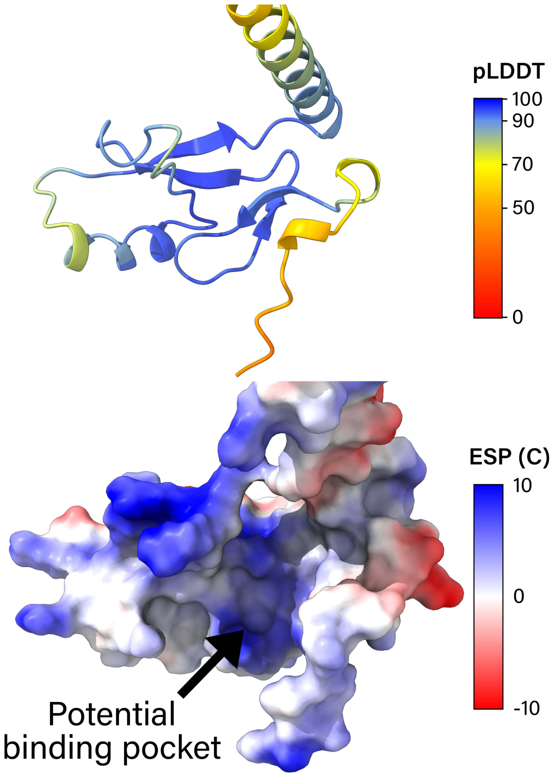

(b)

Figure 21: (a) At the top, a cartoon representation of the binding pocket in the ZZ-domain of HERC2 (PDB: 6WW4). At the bottom, the molecular surface representation of the binding pocket, colored by Coulombic electrostatic potential (ESP). (b) At the top, a cartoon representation of the potential binding pocket in the ZZ-domain of D0LYE7, colored by pLDDT. At the bottom, the molecular surface representation of the potential binding pocket, colored by Coulombic electrostatic potential (ESP).

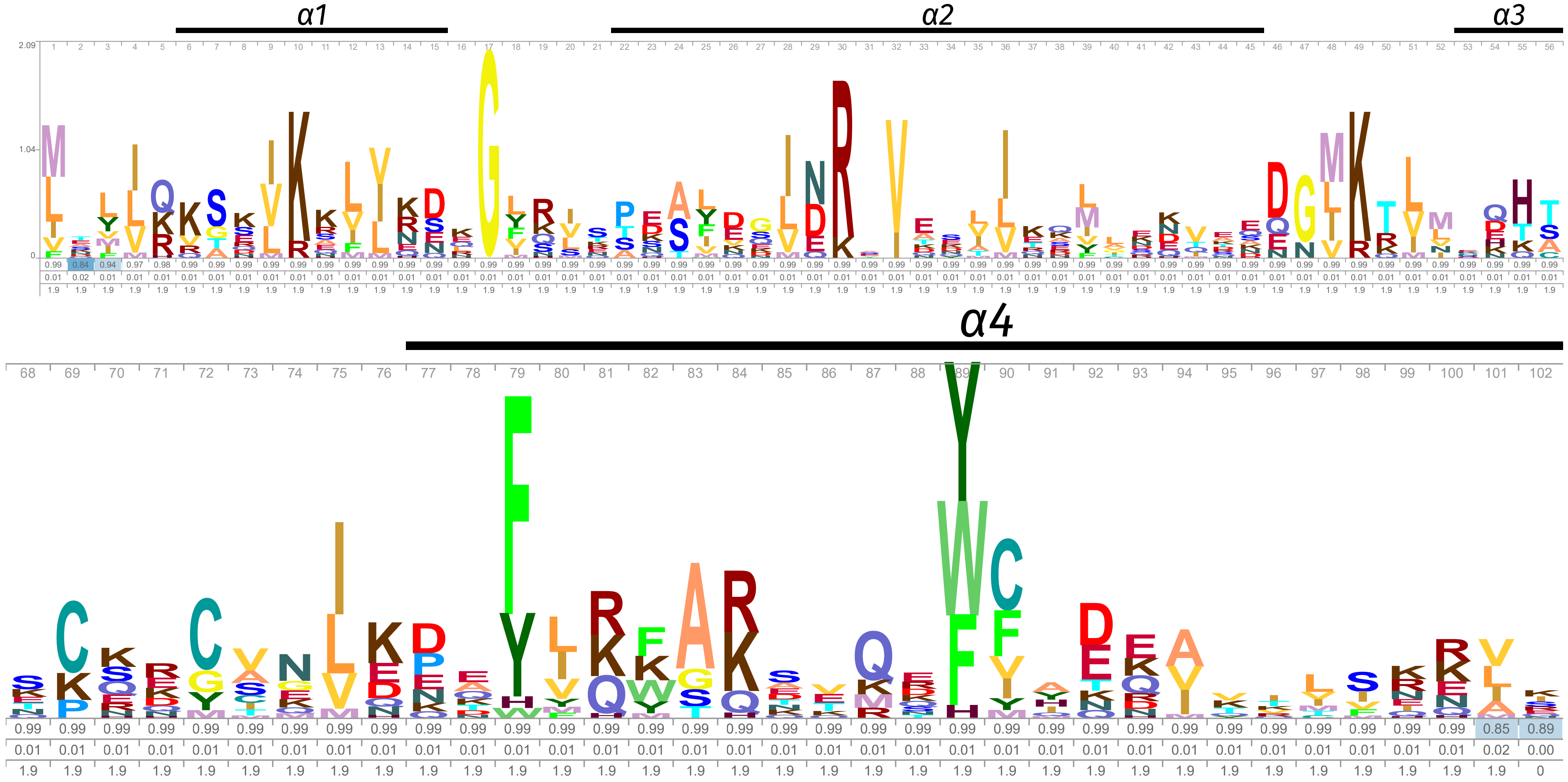

Figure 22: Logo representation of the phage histone HMM profile. Only residues with scores above background frequency are shown. The occupancy probability, insert probability, and insert length values are below the residues. The secondary structure is visualized above the residues.

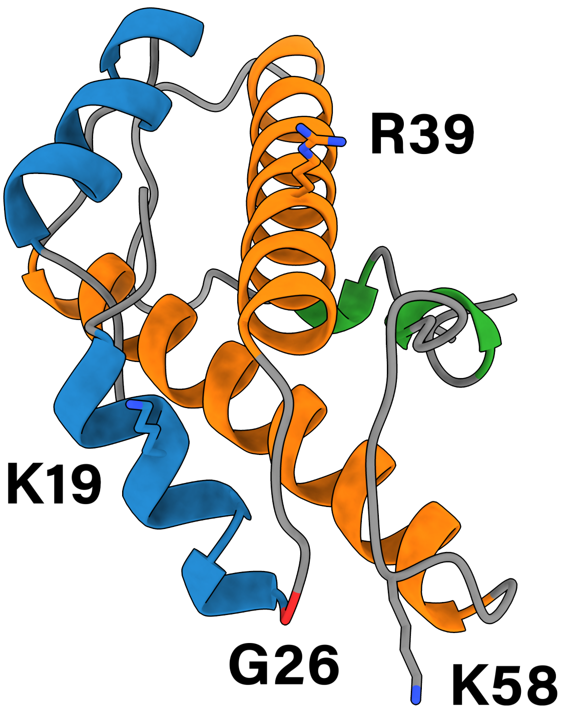

Figure 23: Conserved DNA binding residues K19 and K58, and the conserved G26 and R39 residues of phage histone A0A2E7QIQ9. Residues K19, G26, R39, and K58 correspond to residues K10, G17, R30, and K49 in the HMM logo (Supplementary Fig. 22).

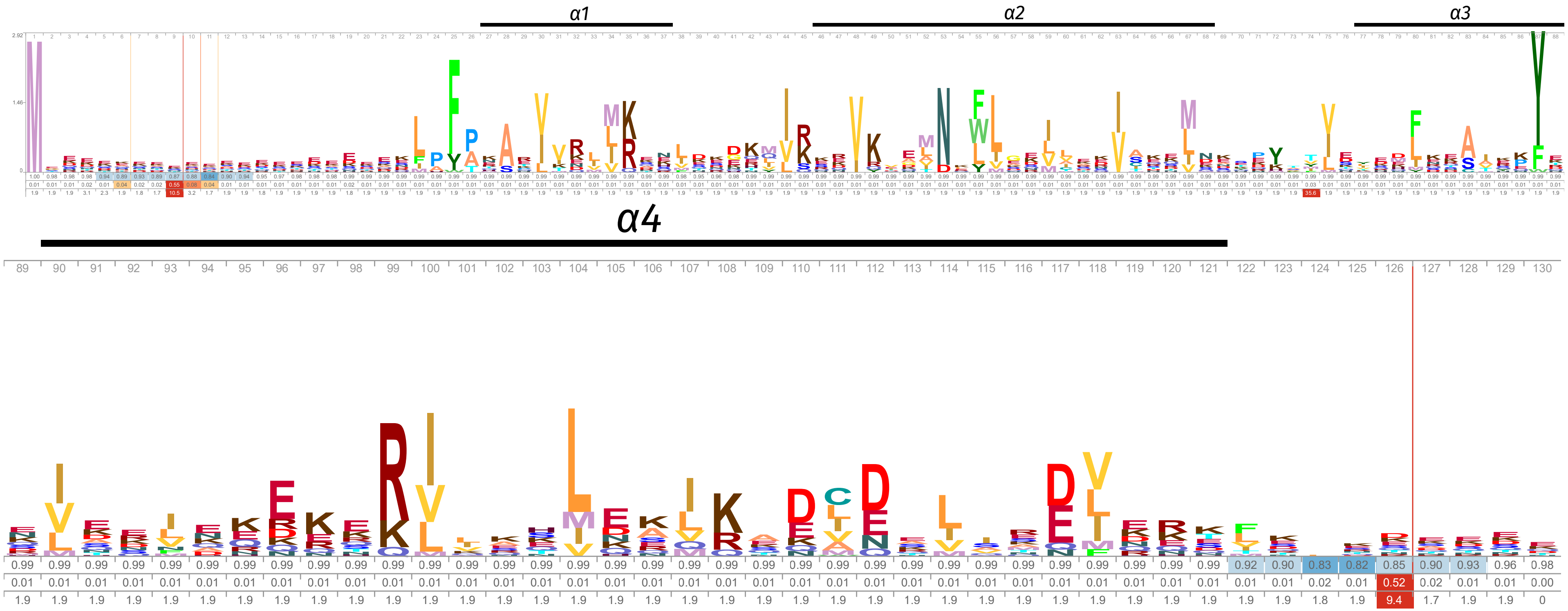

Figure 24: Logo representation of the coiled-coil histone HMM profile. Only residues with scores above background frequency are shown. The occupancy probability, insert probability, and insert length values are below the residues. The secondary structure is visualized above the residues.

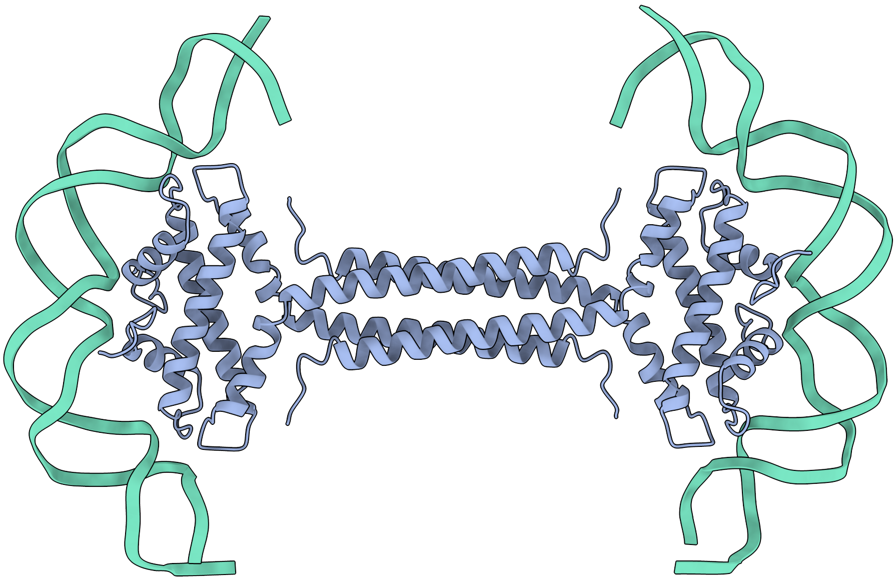

(a)

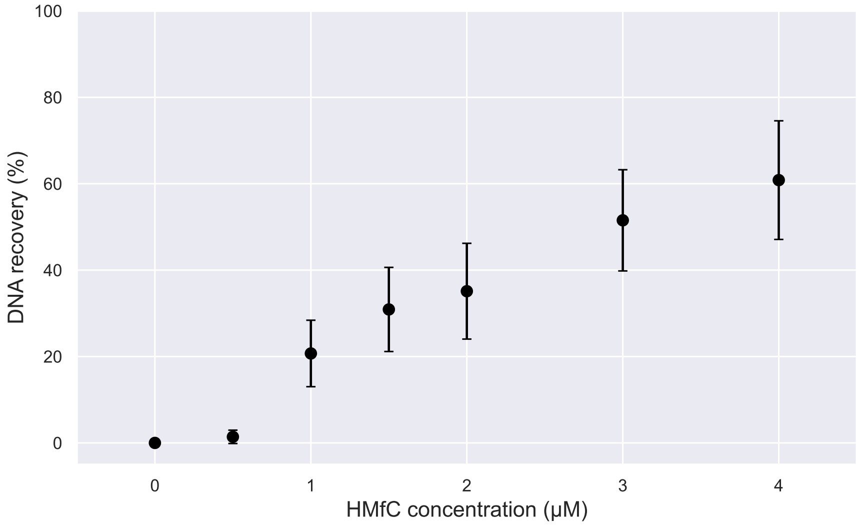

(b)

Figure 25: (a) Our proposed model for how HMfC binds and bridges DNA. (b) DNA bridging assay with HMfC. DNA bridging activity is represented on the y-axis as the percentage of radioactively labeled DNA recovered after pulling-down the magnetic-bead immobilized DNA. Points represent the mean values of three independent measurements. One standard deviation is visualized as error bars.

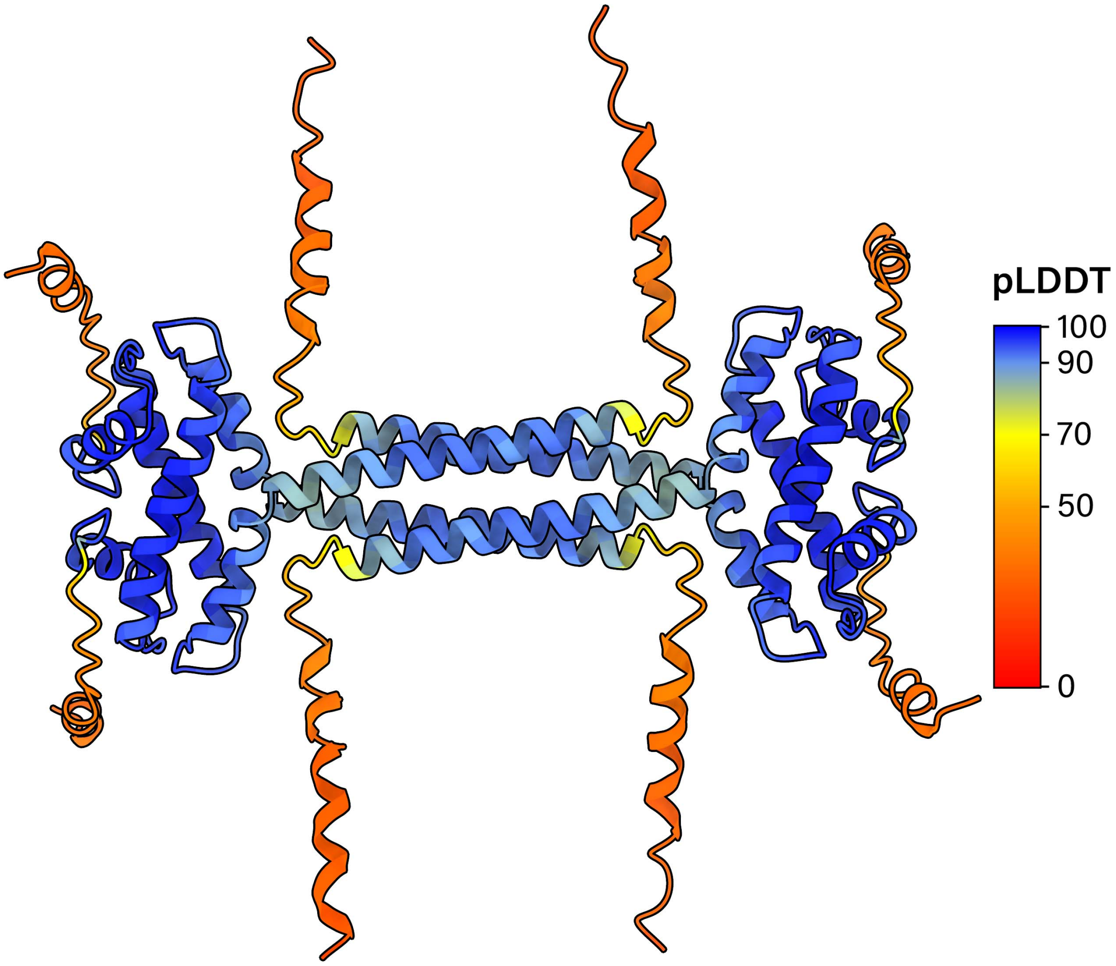

(a)

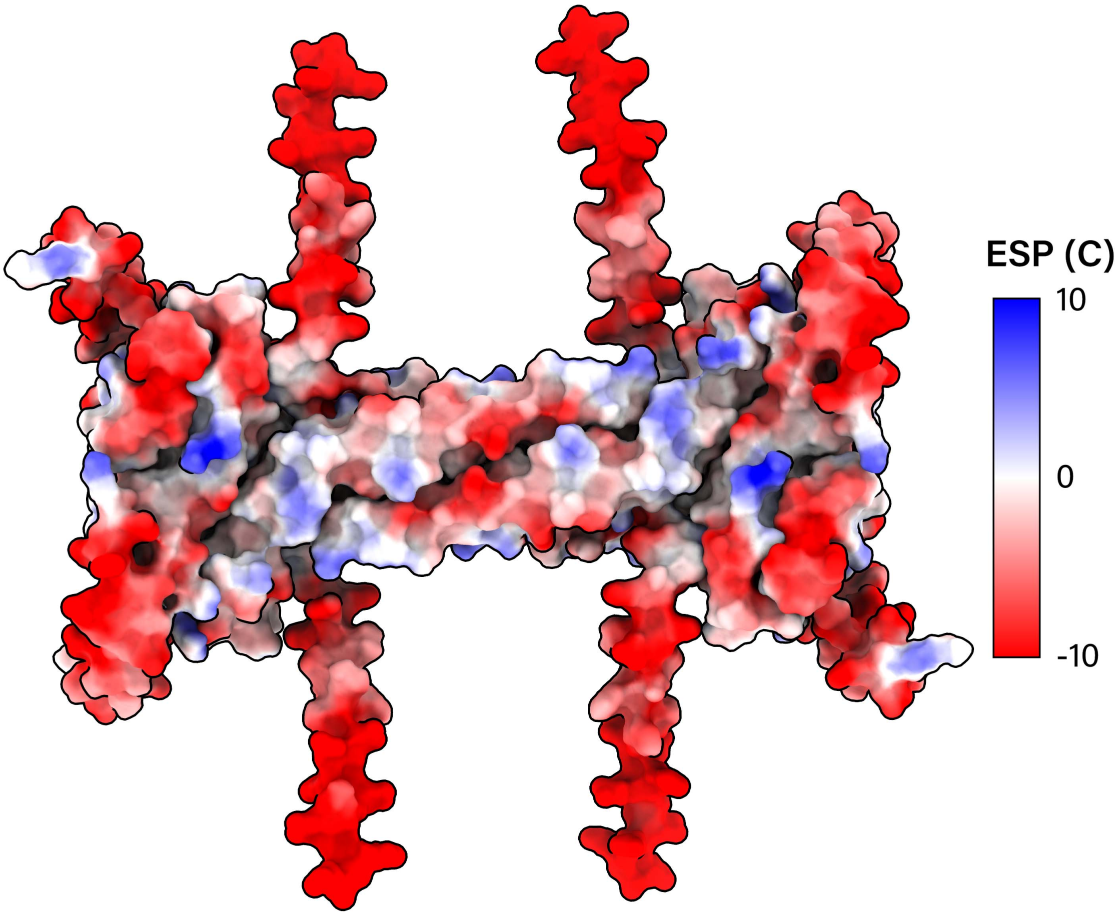

(b)

Figure 26: (a) The homotetramer of A5UK87 as predicted by AlphaFold2. A5UK87 is a coiled-coil histone from *Methanobrevibacter smithii* which contains N- and C-terminal tails which are common in *Methanomada* and *DPANN*. Each residue is colored by its pLDDT value. Values below 50 indicate a prediction of disorder. (b) The molecular surface of A5UK87's tetramer prediction, colored by Coulombic electrostatic potential (ESP).

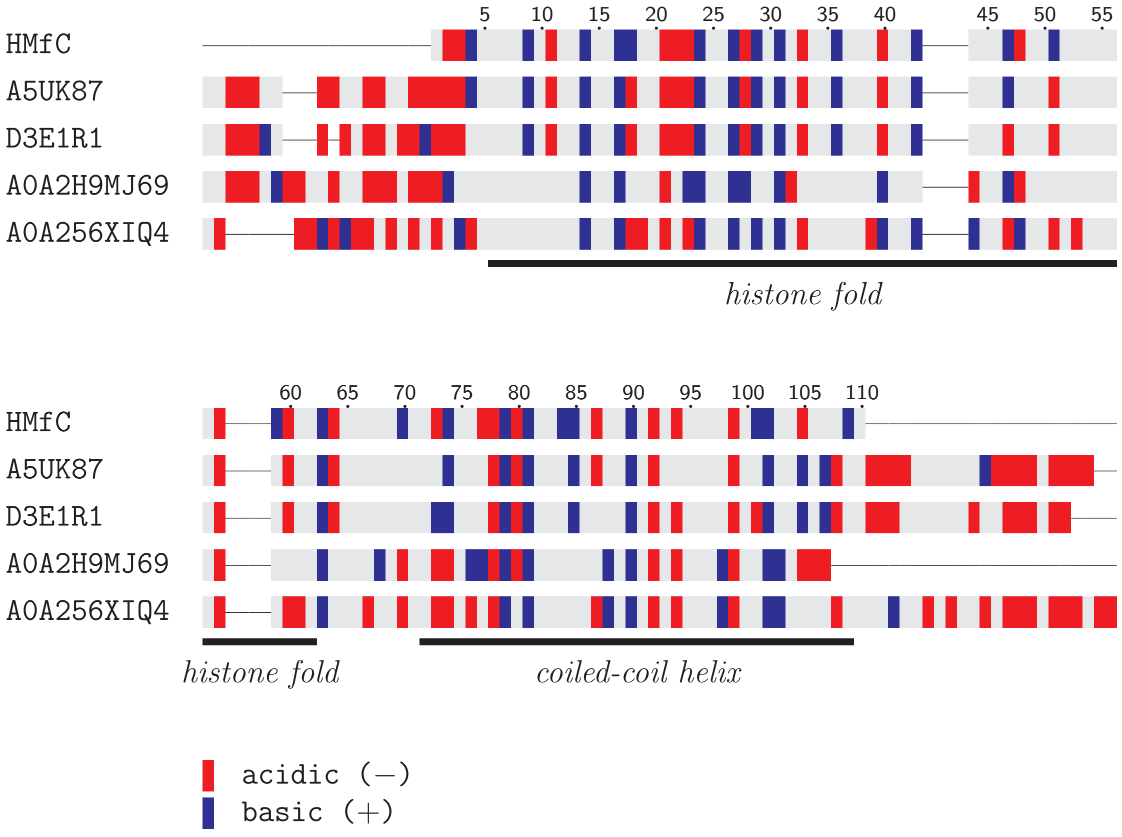

Figure 27: The charge distribution of some coiled-coil histones. The secondary structures are indicated below the sequences. The alignment was made with TCoffee using default parameters. The UniProt accession number for HMfC is E3GZL0.

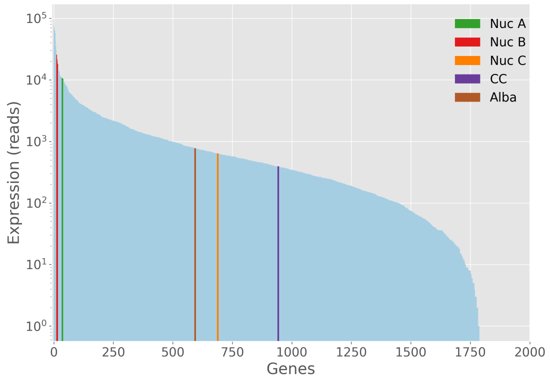

Figure 28: Transcriptome plots of *Methanobrevibacter smithii* PS. mRNA transcriptome data from Hansen et al.. Genes are ranked from highest expression (at x=0) to lowest expression. Only the top 2000 expressed genes are visualized. The y-axis is on a base 10 logarithmic scale. Legend abbreviations are Nuc A:nucleosomal histone A, CC:coiled-coil histone.

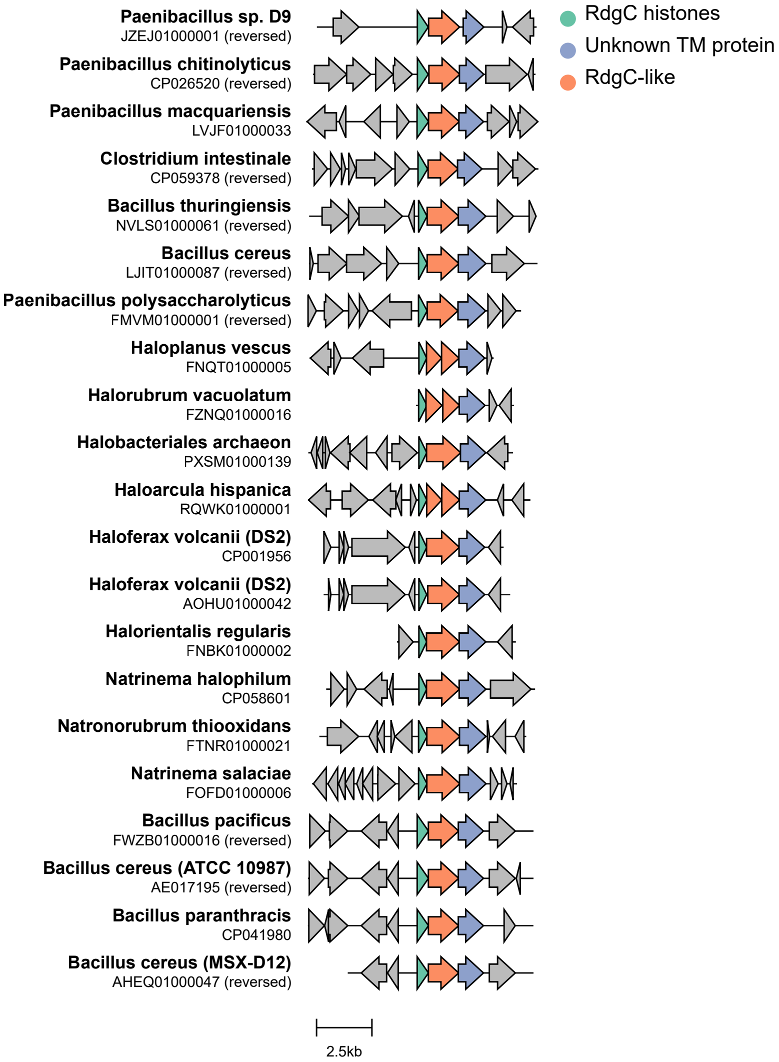

Figure 29: Gene cluster comparison of bacterial genomes which contain the RdgC histone. The organism and its genome ID are noted on the left. Some organisms, such as *Haloplanus vescus*, contain two copies of the RdgC-like protein. In these cases, the RdgC-like protein is split into two parts, each part being encoded by a separate gene.

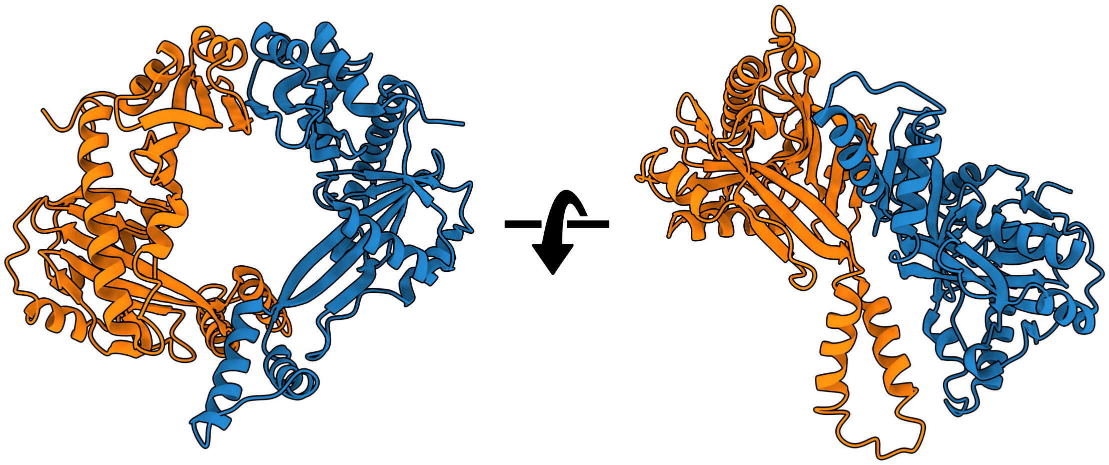

(a)

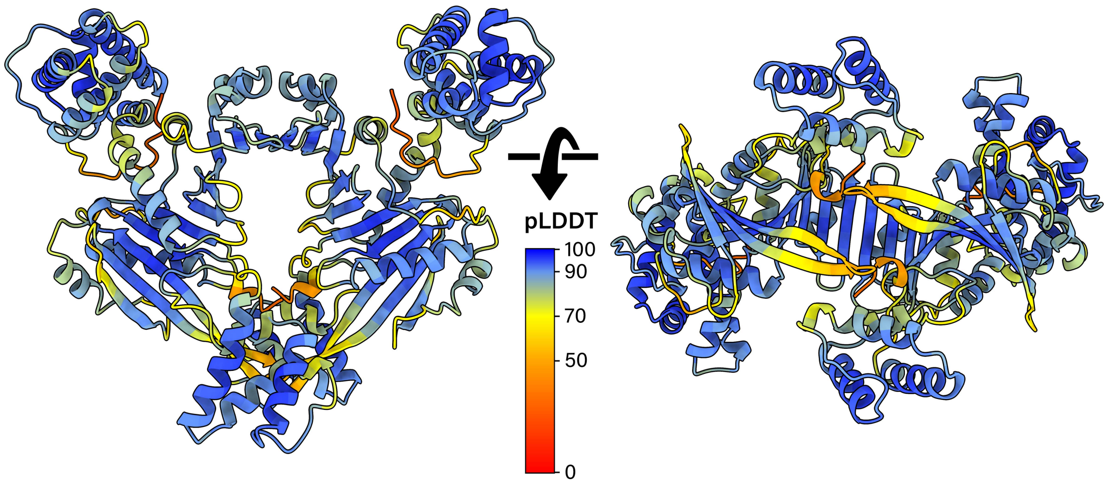

(b)

Figure 30: (a) The RdgC dimer from *Escherichia coli* strain K12 (PDB: 2OWL). (b) The homodimer of RdgC-like protein Q74P82 from *Bacillus cereus* strain ATCC 10987 as predicted by AlphaFold2. Each residue is colored by its pLDDT value.

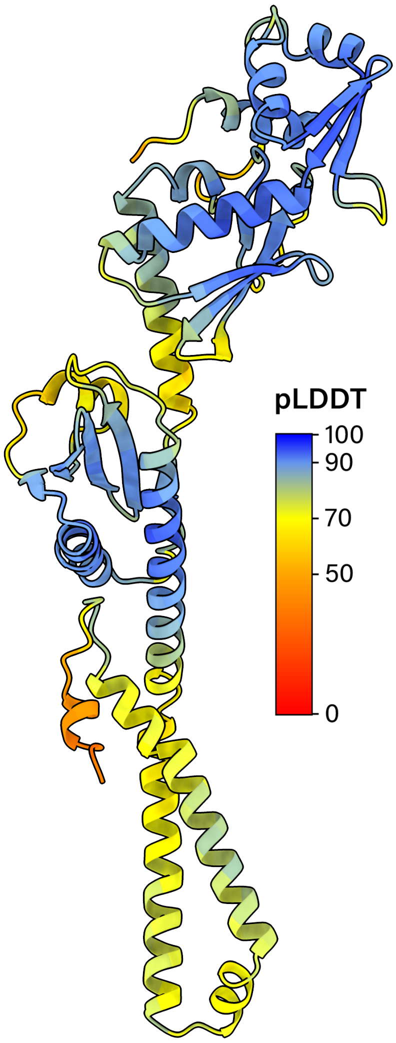
pLDDT
100
90
70
50
0

(a)

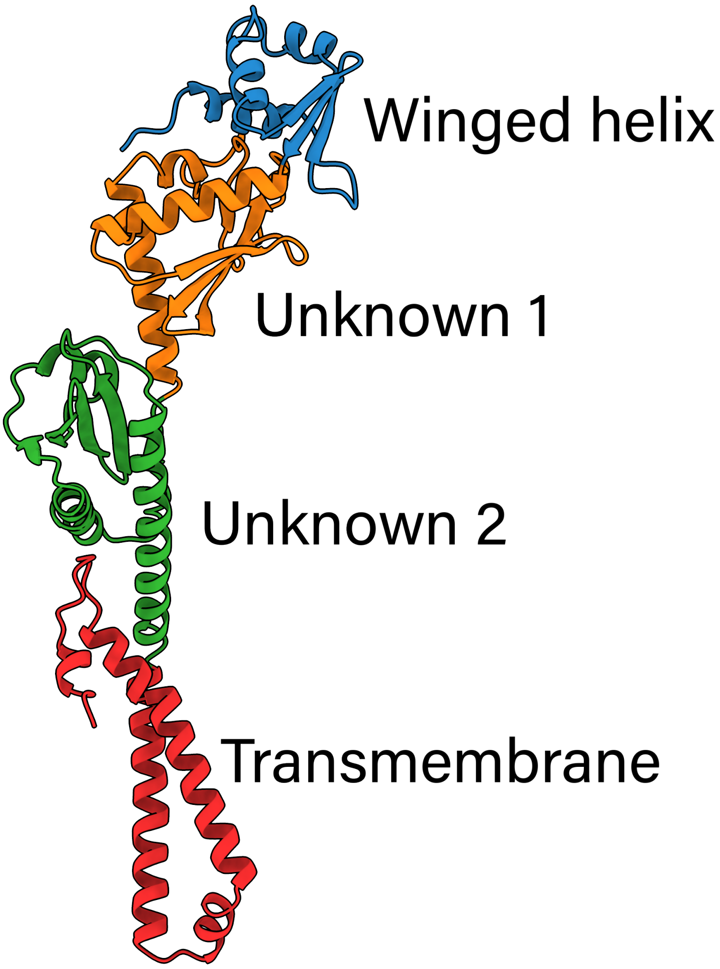
Winged helix
Unknown 1
Unknown 2
Transmembrane

(b)

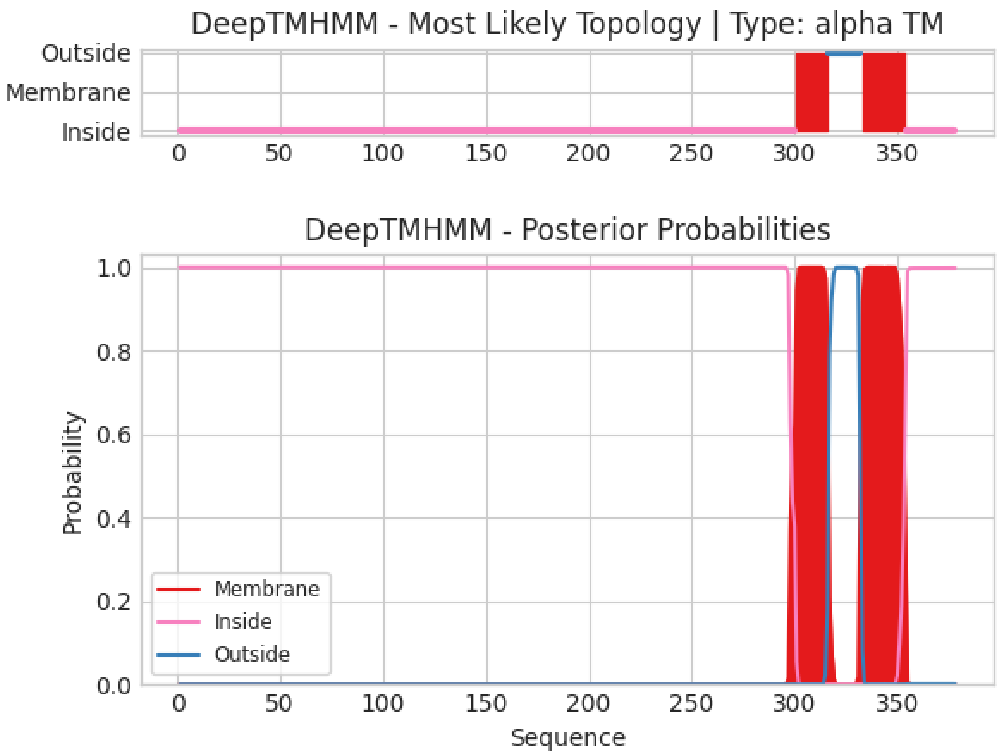

(c)

Figure 31: (a) The monomer of the transmembrane protein Q74P83 from *Bacillus cereus* strain ATCC 10987 as predicted by AlphaFold2 (retrieved from https://alphafold.ebi.ac.uk). Each residue is colored by its pLDDT value. (b) The four different domains that make up Q74P83, each colored separately. YdjF (P77721), YBR085C-A (O43137), and profilin (Q58NA1) are the closest homologs to the winged-helix domain, unknown domain 1, and unknown domain 2 respectively. However, sequence identity is below 30% in all three cases. (c) Prediction of Q74P83's transmembrane probability and its topology by DeepTMHMM.

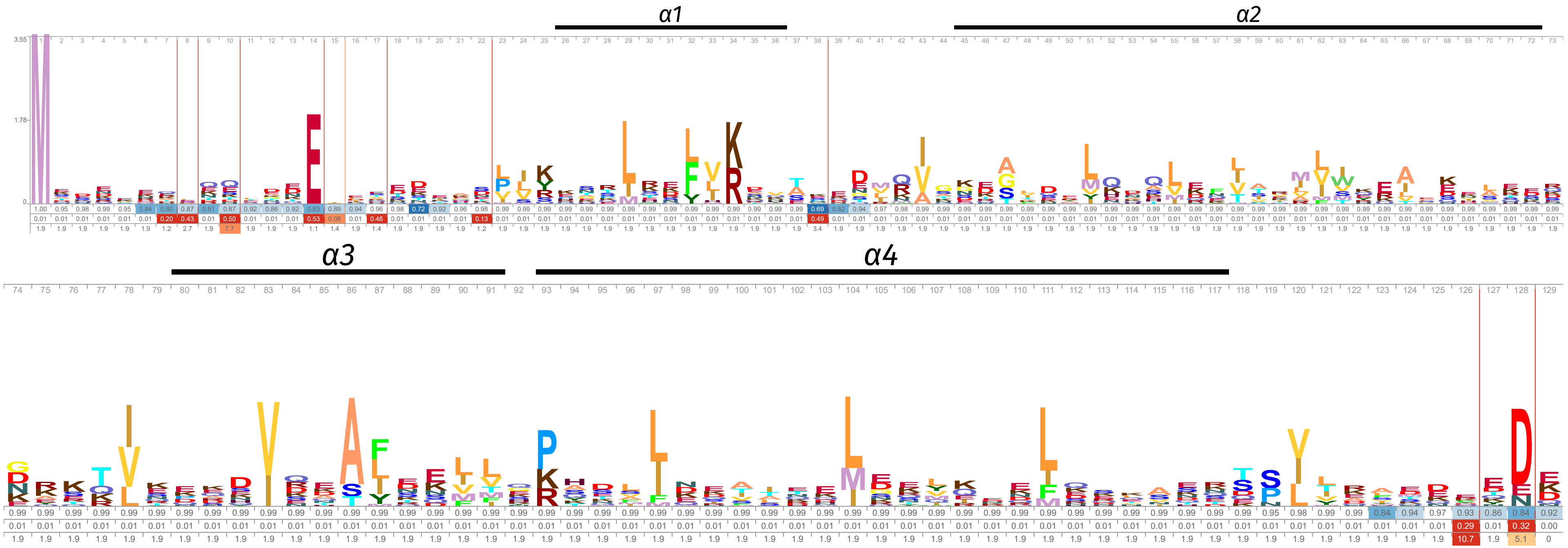

Figure 32: Logo representation of the RdgC histone HMM profile. Only residues with scores above background frequency are shown. The occupancy probability, insert probability, and insert length values are below the residues. The secondary structure is visualized above the residues.

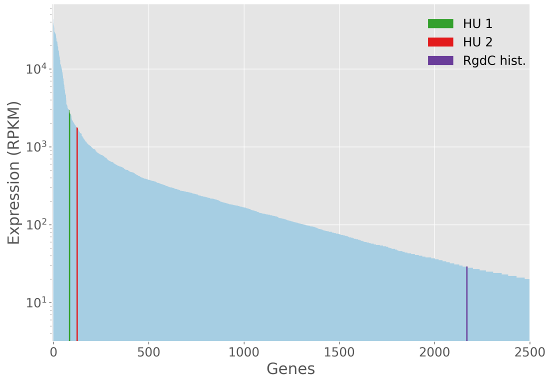

Figure 33: Transcriptome plots of *Bacillus cereus* strain ATCC 10987. mRNA transcriptome data from Kristoffersen et al.. Genes are ranked from highest expression (at x=0) to lowest expression. Only the top 2500 expressed genes are visualized. The y-axis is on a base 10 logarithmic scale. Legend abbreviations are RdgC hist.:RdgC histone.

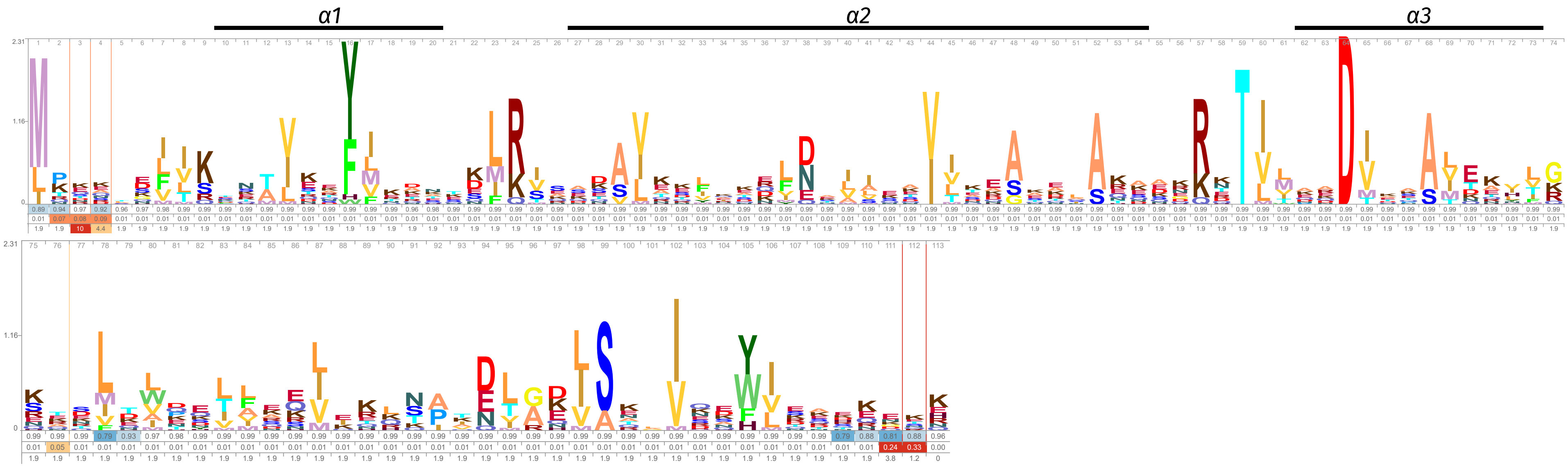

Figure 34: Logo representation of the IHF histone HMM profile. Only residues with scores above background frequency are shown. The occupancy probability, insert probability, and insert length values are below the residues. The secondary structure is visualized above the residues.

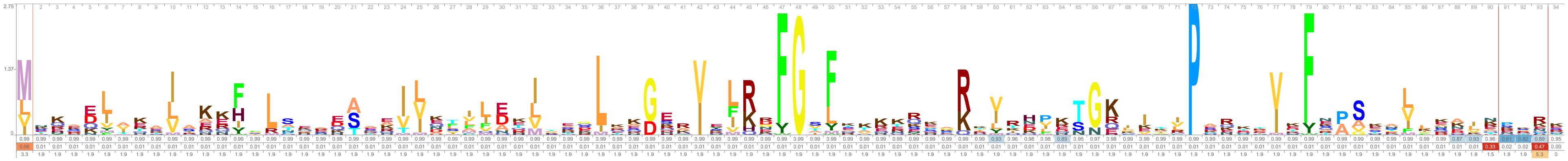

Figure 35: Logo representation of the IHF-like HMM profile. Only residues with scores above background frequency are shown. The occupancy probability, insert probability, and insert length values are below the residues. The secondary structure is visualized above the residues.

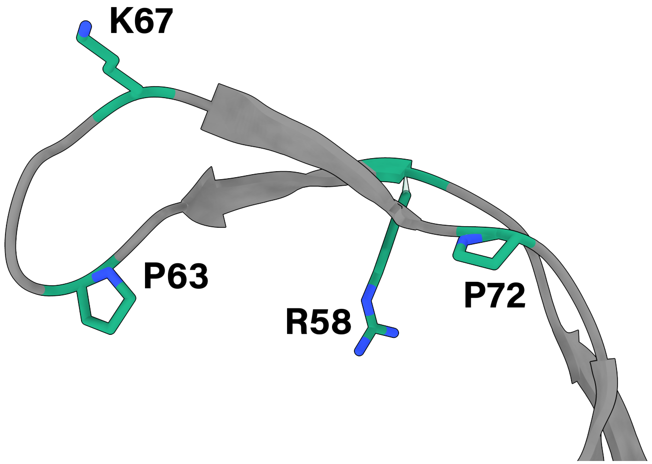

Figure 36: The beta arm of IHF-like A0A358AGI6 as predicted by AlphaFold2. Conserved DNA binding residues R58, K67, P72 and intercalator residue P63 are shown in green. In some IHF-like proteins, P63 is replaced by a different intercalating hydrophobic residue (valine, leucine, and isoleucine)

```
Q6MRM1/1-64    MA......................................     2
D0LYE7/1-180   MSTSNDSAQDPAERSFWKRCSSCKSEIAFGALHFVCSVST    40

Q6MRM1/1-64    ........................................     2
D0LYE7/1-180   CNRKRTGMKFCSVPCWEVHLPMMRHREAYAEEERAPSKDA    80

Q6MRM1/1-64    ..................................EVLVVT     8
D0LYE7/1-180   WLREQRAEAEAQESSARRRTVVAPAAKSSTSMDDEILVVA   120
                                                  histone fold

Q6MRM1/1-64    SKVKKLIKEKGQMNTSAETIDVLSKAIEQLCLKGVESAKA    48
D0LYE7/1-180   SKLKKYIRERSGMNTSDTLLPVLSAHLRVLANQAIRHAAQ   160
                                histone fold

Q6MRM1/1-64    DGRKTVMARDIVIDH....L    64
D0LYE7/1-180   DGRKTVMDRDLAMPSQRDED   180
                  histone fold
```

X non-conserved
X similar
X $\geq$ 50% conserved
X all match

Figure 37: Multiple sequence alignment of histones Q6MRM1 and D0LYZ1 from *Bdellovibrio bacteriovorus* HD100, and *Haliangium ochraceum* SMP-2 respectively. The alignment was made with TCoffee using default parameters.

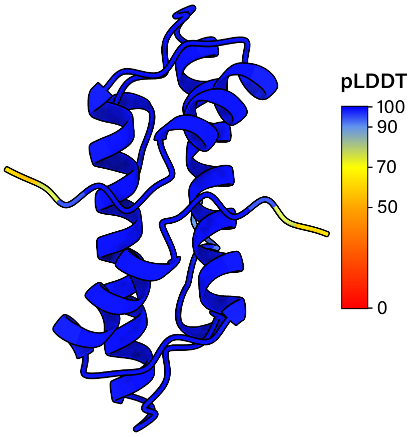

(a)

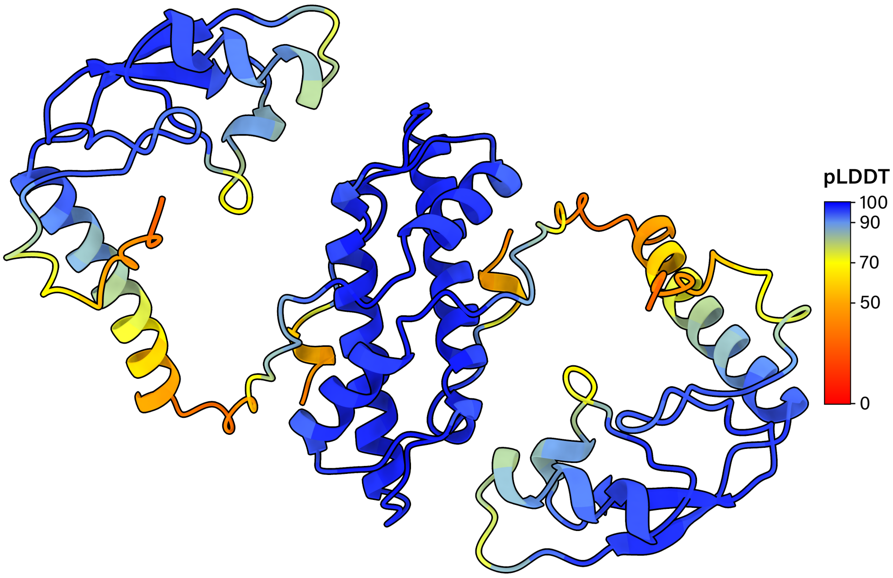

(b)

Figure 38: (a) The homodimer of bacterial dimer histone Q6MRM1 from *Bdellovibrio bacteriovorus* HD100 as predicted by AlphaFold2. Each residue is colored by its pLDDT value. (b) The homodimer of ZZ histone D0LYE7 from *Haliangium ochraceum* SMP-2 as predicted by AlphaFold2. Each residue is colored by its pLDDT value.

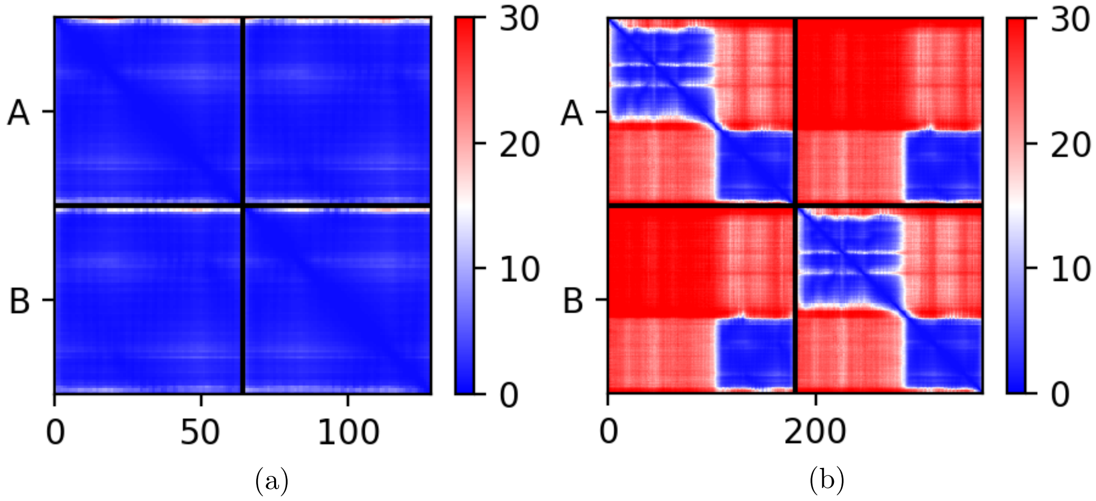

Figure 39: Predicted aligned error plots for the (a) Q6MRM1 homodimer and (b) D0LYE7 homodimer predictions.

### References

[1] John Jumper, Richard Evans, Alexander Pritzel, Tim Green, Michael Figurnov, Olaf Ronneberger, Kathryn Tunyasuvunakool, Russ Bates, Augustin Žídek, Anna Potapenko, Alex Bridgland, Clemens Meyer, Simon A.A. Kohl, Andrew J. Ballard, Andrew Cowie, Bernardino Romera-Paredes, Stanislav Nikolov, Rishub Jain, Jonas Adler, Trevor Back, Stig Petersen, David Reiman, Ellen Clancy, Michal Zielinski, Martin Steinegger, Michalina Pacholska, Tamas Berghammer, Sebastian Bodenstein, David Silver, Oriol Vinyals, Andrew W. Senior, Koray Kavukcuoglu, Pushmeet Kohli, and Demis Hassabis. Highly accurate protein structure prediction with AlphaFold. *Nature*, 596(7873):583–589, 2021.

[2] Richard Evans, Michael O'Neill, Alexander Pritzel, Natasha Antropova, Andrew Senior, Tim Green, Augustin Žídek, Russ Bates, Sam Blackwell, Jason Yim, Olaf Ronneberger, Sebastian Bodenstein, Michal Zielinski, Alex Bridgland, Anna Potapenko, Andrew Cowie, Kathryn Tunyasuvunakool, Rishub Jain, Ellen Clancy, Pushmeet Kohli, John Jumper, and Demis Hassabis. Protein complex prediction with alphafold-multimer. *bioRxiv*, 2021.

[3] Joana Pereira, Adam J. Simpkin, Marcus D. Hartmann, Daniel J. Rigden, Ronan M. Keegan, and Andrei N. Lupas. High-accuracy protein structure prediction in CASP14. *Proteins: Structure, Function, and Bioinformatics*, 89(12):1687–1699, 2021.

[4] Arne Elofsson. Progress at protein structure prediction, as seen in CASP15. *Current Opinion in Structural Biology*, 80:102594, June 2023.

[5] Kathryn Tunyasuvunakool, Jonas Adler, Zachary Wu, Tim Green, Michal Zielinski, Augustin Žídek, Alex Bridgland, Andrew Cowie, Clemens Meyer, Agata Laydon, Sameer Velankar, Gerard J. Kleywegt, Alex Bateman, Richard Evans, Alexander Pritzel, Michael Figurnov, Olaf Ronneberger, Russ Bates, Simon A. A. Kohl, Anna Potapenko, Andrew J. Ballard, Bernardino Romera-Paredes, Stanislav Nikolov, Rishub Jain, Ellen Clancy, David Reiman, Stig Petersen, Andrew W. Senior, Koray Kavukcuoglu, Ewan Birney, Pushmeet Kohli, John Jumper, and Demis Hassabis. Highly accurate protein structure prediction for the human proteome. *Nature*, 596(7873):590–596, August 2021.

[6] Mehmet Akdel, Douglas E. V. Pires, Eduard Porta Pardo, Jürgen Jänes, Arthur O. Zalevsky, Bálint Mészáros, Patrick Bryant, Lydia L. Good, Roman A. Laskowski, Gabriele Pozzati, Aditi Shenoy, Wensi Zhu, Petras Kundrotas, Victoria Ruiz Serra, Carlos H. M. Rodrigues, Alistair S. Dunham,

David Burke, Neera Borkakoti, Sameer Velankar, Adam Frost, Jérôme Basquin, Kresten Lindorff-Larsen, Alex Bateman, Andrey V. Kajava, Alfonso Valencia, Sergey Ovchinnikov, Janani Durairaj, David B. Ascher, Janet M. Thornton, Norman E. Davey, Amelie Stein, Arne Elofsson, Tristan I. Croll, and Pedro Beltrao. A structural biology community assessment of AlphaFold2 applications. *Nature Structural & Molecular Biology*, 29(11):1056–1067, November 2022.
